# Sequencing the gaps: dark genomic regions persist in CHM13 despite long-read advances

**DOI:** 10.1101/2025.05.23.655776

**Authors:** Mark E. Wadsworth, Madeline L. Page, Bernardo Aguzzoli Heberle, Justin B. Miller, Cody Steely, Mark T. W. Ebbert

## Abstract

Comprehensive genomic analysis is essential for advancing our understanding of human genetics and disease. However, short-read sequencing technologies are inherently limited in their ability to resolve highly repetitive, structurally complex, and low-mappability genomic regions, previously coined as “dark” regions. Long-read sequencing technologies, such as PacBio and Oxford Nanopore Technologies (ONT), offer improved resolution of these regions, yet they are not perfect. With the advent of the new Telomere-to-Telomere (T2T) CHM13 reference genome, exploring its effect on dark regions is prudent. In this study, we systematically analyze dark regions across four human genome references—HG19, HG38 (with and without alternate contigs), and CHM13—using both short- and long-read sequencing data. We found that dark regions increase as the reference becomes more complete, especially dark-by-MAPQ regions, but that long-read sequencing significantly reduces the number of dark regions in the genome, particularly within gene bodies. However, we identify potential alignment challenges in long-read data, such as centromeric regions. These findings highlight the importance of both reference genome selection and sequencing technology choice in achieving a truly comprehensive genomic analysis.

## Introduction

The ultimate goal of human genomics research is to improve disease diagnostics and treatments. To this end, performing a comprehensive and personalized analysis on an individual’s genome, transcriptome, and epigenome, in combination with other factors (e.g., environmental factors) is essential. Specifically, a “complete” analysis at the DNA level should, at minimum, capture all DNA variants (small and structural) that the individual carries, and interpret and predict the downstream implications of these variants. Unfortunately, many gaps remain, but major technological and scientific advances in the past decade have provided significant gains toward that goal. Specifically, short-read sequencing became widely accessible in the early 2010’s, propelling our understanding of the human genome and transcriptome to a level that was difficult to imagine only a decade before. While short-read sequencing has been a major boon to genomics research, many genomic regions remained inaccessible (i.e., dark or camouflaged) and structural variants could not be accurately resolved because of the inherent limitations of short read lengths [1–3].

In 2019, we systematically characterized the long-known, but poorly appreciated issue surrounding the “dark” genome—regions of the genome that cannot be accurately resolved and are thus entirely overlooked [1]. Specifically, using short-read sequencing we characterized two basic forms of “dark” regions, including: (1) regions that are dark-by-depth, where few or no sequencing reads are present in the alignment; and (2) dark-by-mapping-quality (dark-by-MAPQ) where the region contains aligned reads, but the reads do not align uniquely to that region. In both cases, any variants within the region are completely overlooked using standard short-read sequencing and downstream analyses because of ambiguous alignments [1–3]. We further demonstrated the breadth of this issue and how it affects genes already known to be involved in human disease. We termed genes containing dark-by-MAPQ regions because of either full or partial genomic duplications as “camouflaged” genes [1]. In all, we identified >6000 gene bodies where some portion of the gene’s sequence is “dark” using standard short-read sequencing approaches, and 2128 were ≥ 5% dark [1]. Dark and camouflaged genes make it difficult (if not impossible) to perform a complete analysis of an individual’s genome, including for both small and large DNA variants [1]. We also showed that long-read sequencing resolves most dark and camouflaged regions overlooked by short-read sequencing [1]. More recent work has shown that long reads also characterize and quantify individual RNA isoform expression [4–7], ultimately bringing us closer to the reality of a comprehensive and personalized genomic analysis.

A complete and accurate representation of the human genome is essential to understanding its complexity, but the human genome has only recently been completed. Leveraging long-read sequencing, the Telomere-2-Telomere (T2T) consortium assembled the first complete human genome sequence from a completely homozygous cell line (CHM13) in 2022 [8, 9]. The new T2T CHM13 reference genome added ∼200 megabases to the previously most complete reference genome (GRCh38), predominantly in telomeric, centromeric, and acrocentric chromosomal regions [8].

Though tempting to assume the human genome is finally “complete”, as the efforts and latest results from the Human Pangenome Reference Consortium demonstrate, the human genome is still far from “complete” because no single reference can accurately represent all of humanity [10]; individuals have their own unique combination of not only single-nucleotide variants, but also larger structural DNA variations [11–14]. Thus, being able to perform a complete analysis of an individual’s genome (whether by *de novo* assembly or alignment to a reference genome) is essential to understanding their genetic predisposition for various phenotypes, including those involved in human health and disease [11–13, 15]. Unfortunately, we are still a long way from achieving the goal to perform a truly complete analysis of an individual’s genome, especially since most clinical and research still rely on standard short-read sequencing approaches and pipelines that are known to overlook critical regions of an individual’s genome [1].

To perform a comprehensive analysis, we first need a “comprehensive” reference genome to serve as a baseline, for which CHM13 and the pangenome are the beginning. Counterintuitively, however, the more complete the human reference genome becomes, the more challenging it becomes to properly analyze and interpret an individual’s unique combination of DNA variants because of genomic duplications and ambiguous alignments. In this challenging area of research, we address three important knowledge gaps, herein: (1) how the interaction between sequencing platform choice and reference genome, especially CHM13, affects our ability to assess the “dark” and “camouflaged” regions of the genome; (2) how dark and camouflaged regions affect related short-read sequencing assays beyond standard DNA sequencing (e.g., ChIP-Seq, bisulfite sequencing, etc.) and further prevent a complete genomic analysis using short-read data; and (3) the importance of supplementary alignments in long-read datasets.

## Results

To better understand how the intersection of sequencing length/platform and reference genome affects our ability to accurately resolve dark and camouflaged regions of the genome, we compared dark regions from long-read (PacBio and Oxford Nanopore Technologies) and short-read sequencing data (Illumina 100bp reads and 250bp reads; **Fig. 1a**) across four different human genome references. We compared reference genomes representing a continuum of genome completeness: (1) HG19; (2) HG38 (excluding alternate contigs); (3) HG38 (including alternate contigs); and (4) the Telomere-to-Telomere (T2T) CHM13 v2.0 (**Fig. 1b**). Using the same 20 short-read samples from our original paper (ten 100bp Illumina and ten 250bp Illumina) [1], and ten samples sequenced on both PacBio and ONT (**Fig. 1a,b**), we aligned each sample to the four reference genomes (excluding secondary and supplementary alignments, per our 2019 analysis) and identified dark and camouflaged regions using the Dark Region Finder (**Fig. 1c**) [1]. For clarity, secondary reads are those that map equally well in multiple places in the genome while supplementary reads are chimeric reads, mapping different regions of the same read independently. We then quantified dark-by-depth and dark-by-MAPQ regions across platforms and references genome-wide. Building upon our previous work, we performed a comparison of dark regions between HG38 and T2T’s CHM13 (**Fig. 1di**) [8, 16, 17]. As part of our analyses, we also assessed the effect of dark and camouflage regions in other short-read based sequencing assays, including epigenetic assays like ChIP-Seq and bisulfite sequencing (**Fig. 1dii**).

**Figure 1:**
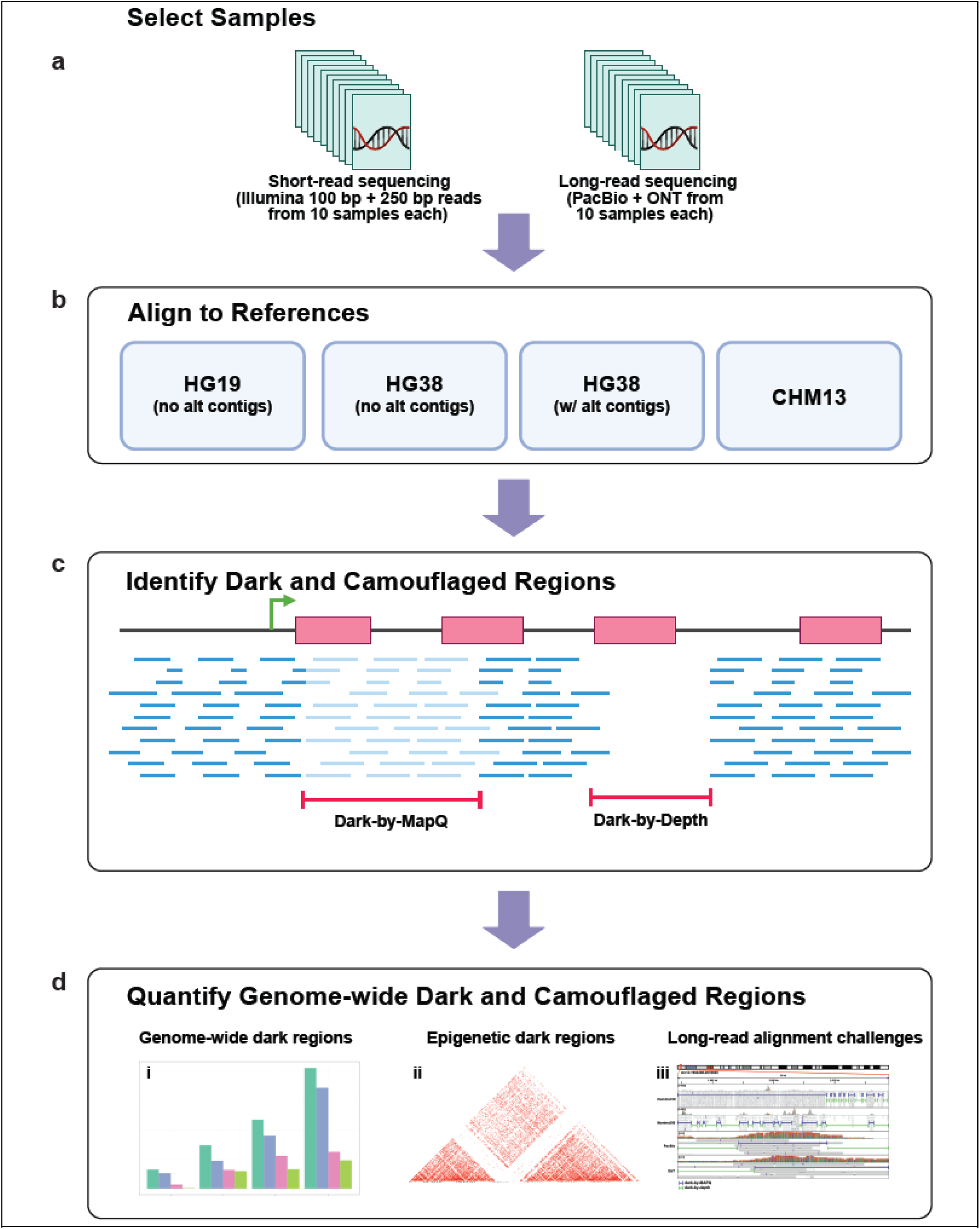
Dark region analytical workflow. **(a)** We obtained 20 samples sequenced using Illumina short-reads (100bp and 250bp) and 10 samples sequenced using PacBio and ONT. **(b)** Using bwa and minimap2 for short and long reads, respectively, we aligned all samples to HG19, HG38 (with and without alternate contigs), and CHM13. **(c)** We then identified dark and camouflaged regions using our Dark Region Finder. **(d) i.** Finally, we characterized and quantified dark regions genome wide, along with their differences between references and sequencing platforms. **ii.** We further assessed how these dark and camouflaged regions affect other short-read based DNA assays (i.e., epigenetic assays). **iii.** We identified important long-read alignment challenges that need to be addressed. Created with BioRender.com.

In our 2019 work, we excluded secondary and supplementary alignments for simplicity. Here, we demonstrate that excluding supplementary reads falsely inflates estimates for dark-by-depth regions in long-read data, and thus demonstrates the importance of supplementary reads. We discovered this because of a region that was dark-by-depth in long reads but resolved in short reads when excluding supplementary reads. Upon deeper investigation, we found that including supplementary reads resolved this issue.

### Number of dark regions increase in CHM13

In our previous work, we characterized and quantified dark and camouflaged regions across the genome using short-read sequencing technologies and assessed how well long reads resolved these regions, when using primary alignments only [1]. Here, we build on that work by including CHM13. After aligning the samples to each of the four reference genomes, we classified dark regions into three main classes: (1) dark-by-depth regions, which have <5x coverage; (2) dark-by-MAPQ regions, in which 90% of the reads covering the region have a mapping quality (MAPQ) less than 10; and (3) camouflaged regions, which are a subset of dark-by-MAPQ regions that are also 98% identical to another genomic region, as calculated by BLAT [18].

We observed several important patterns when comparing genome-wide dark-by-MAPQ bases for each platform across the four reference genomes. As the reference genome became more complete, between HG19 and CHM13, the number of dark-by-MAPQ bases increased. Specifically, there was >6x increase in dark-by-MAPQ bases in CHM13 compared to HG19 for Illumuna100 reads (**Fig. 2a**; **Table 1**). In CHM13, there are 3.3x more dark-by-MAPQ bases in short-read data compared to long-reads (**Fig. 2a**; **Table 1**) [23,24,35]. Specifically, Illumina100 reads had 206,689,498 dark-by-MAPQ bases when aligned to CHM13, while PacBio and ONT had 78,273,227 (resolved 62.1%) and 62,833,689 (resolved 69.6%), respectively. Upon initial inspection, the number of dark-by-depth bases appears to be enriched in long-read technologies (**Fig. 2b**), but we show later that this enrichment is artificially inflated in ONT because we omitted supplementary reads in our 2019 work and in most analyses in this work. When comparing within gene bodies (i.e. not genome wide), PacBio and ONT had 6,071,156 and 1,802,133 dark-by-MAPQ bases, respectively, showing that within gene bodies, ONT resolved 3.4 times more bases than PacBio (**Fig. 2c,d**).

**Figure 2:**
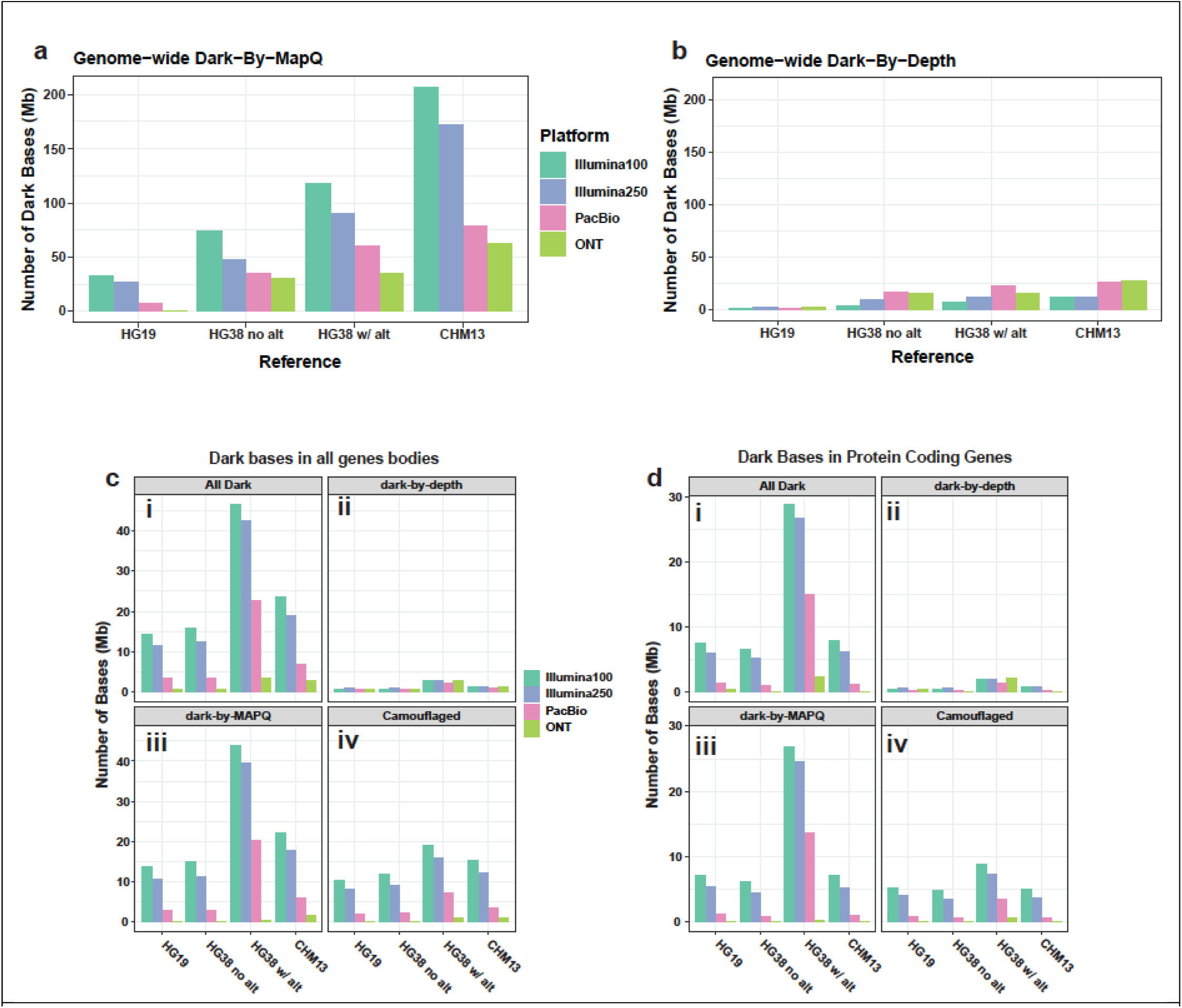
Dark-by-MAPQ dark bases predominate. **(a)** As the genome becomes more complete, short-read data exhibits increasing dark-by-MAPQ nucleotides while long reads plateau. Green and pink arrows show general trends for Illumina100 and PacBio, respectively. **(b)** While millions of bases are obscured by dark-by-depth regions, they pale in comparison to dark-by-MAPQ **(c)** ONT has the least number of dark bases within genes in all types of dark regions **(i)**, dark-by-depth **(ii)**, dark-by-MAPQ **(iii)**, and camouflaged regions **(iv)** across all references. HG38 with alternates has the highest amount of dark bases. **(d)** ONT has the least number of dark bases within protein coding genes in all types of dark regions **(i)**, dark-by-depth **(ii)**, dark-by-MAPQ **(iii)**, and camouflaged regions **(iv)** across all references. HG38 with alternates has the worst amount of dark bases.

**Table 1:** Dark region statistics reveal long-read dark regions are fewer in number but span larger regions. For all four sequencing technologies and all four reference genomes, we compared the number of dark bases, regions, average bases per region, and median number of bases per region for dark-by-depth, dark-by-MAPQ, and the combination of those two. We found that short-read sequencing data has more, smaller dark regions, while long-read sequencing has fewer, larger dark regions.

| Platform | Reference | Dark-by-depth |  |  |  | Dark-by-MAPQ |  |  |  | All Dark |  |  |  |
| --- | --- | --- | --- | --- | --- | --- | --- | --- | --- | --- | --- | --- | --- |
|  |  | Number of Bases | Number of Regions | Mean Bases per Region | Median Bases per Region | Number of Bases | Number of Regions | Mean Bases per Region | Median Bases per Region | Number of Bases | Number of Regions | Mean Bases per Region | Median Bases per Region |
| Illumina100 | HG19 | 1,167,989 | 11,988 | 97.43 | 57 | 32,837,051 | 51,403 | 638.82 | 137 | 34,005,040 | 58,330 | 582.98 | 115 |
| Illumina250 | HG19 | 1,832,263 | 11,219 | 163.32 | 102 | 26,229,311 | 19,364 | 1,354.54 | 583 | 28,061,574 | 25,971 | 1,080.50 | 338 |
| PacBio | HG19 | 1,319,120 | 324 | 4,071.36 | 1,801 | 7,460,817 | 290 | 25,726.96 | 10,843.50 | 8,779,937 | 502 | 17,489.91 | 3,337 |
| ONT | HG19 | 1,464,751 | 195 | 7,511.54 | 2,176 | 691,318 | 24 | 28,804.90 | 1,805.50 | 2,156,069 | 211 | 10,218.34 | 2,176 |
| Illumina100 | HG38 no Alt | 3,721,688 | 40,806 | 91.2 | 55 | 73,843,911 | 105,606 | 699.24 | 196 | 77,565,599 | 96,787 | 801.41 | 158 |
| Illumina250 | HG38 no Alt | 9,163,182 | 37,231 | 246.12 | 133 | 47,018,216 | 53,262 | 882.77 | 437 | 56,181,398 | 63,748 | 881.3 | 347 |
| PacBio | HG38 no Alt | 16,636,630 | 1,141 | 14,580.74 | 3,749 | 35,215,658 | 1,257 | 28,015.64 | 8,924 | 51,852,288 | 1,281 | 40,477.98 | 5,005 |
| ONT | HG38 no Alt | 14,664,911 | 685 | 21,408.63 | 3,114 | 30,564,861 | 569 | 53,716.80 | 1,286 | 45,229,772 | 883 | 51,222.84 | 2,887 |
| Illumina100 | HG38 Alt | 6,914,302 | 54,812 | 126.15 | 61 | 117,906,741 | 128,622 | 916.69 | 186 | 124,821,043 | 120,689 | 1,034.24 | 163 |
| Illumina250 | HG38 Alt | 12,134,914 | 45,093 | 246.12 | 138 | 90,036,424 | 64,821 | 1,389.01 | 472 | 102,171,338 | 74,705 | 1,367.66 | 383 |
| PacBio | HG38 Alt | 21,763,221 | 1,326 | 16,412.69 | 4,910.50 | 60,183,222 | 1,781 | 33,791.81 | 10,859 | 81,946,443 | 1,859 | 44,080.93 | 6,965 |
| ONT | HG38 Alt | 14,708,016 | 687 | 21,409.05 | 3,083 | 34,475,006 | 620 | 55,604.85 | 1,143.50 | 49,183,022 | 928 | 52,998.95 | 2606.5 |
| Illumina100 | CHM13 | 11,269,696 | 97,770 | 115.27 | 68 | 206,689,498 | 236,580 | 873.66 | 198 | 217,959,194 | 208,620 | 1,044.77 | 169 |
| Illumina250 | CHM13 | 11,781,153 | 55,512 | 212.23 | 132 | 171,496,781 | 107,573 | 1,594.24 | 754 | 183,277,934 | 102,599 | 1786.35 | 600 |
| PacBio | CHM13 | 25,762,200 | 2,091 | 12,320.52 | 4,707 | 78,273,227 | 4,094 | 19,119.01 | 5,495 | 104,035,427 | 3,397 | 30,625.98 | 3,435 |
| ONT | CHM13 | 26,747,722 | 1,114 | 24,010.52 | 1,824.50 | 62,833,689 | 1,869 | 33,618.88 | 1,266 | 89,581,411 | 1,551 | 57,757.20 | 1,396 |

### ONT outperforms each technology for resolving dark regions within gene bodies, including in the CHM13 reference genome in primary alignments

After comparing the dark bases genome-wide, we specifically compared dark bases within gene bodies, which was a major focus of our previous work using primary alignments, only [1]. Here, we found that HG38 with alternate contigs had 2-3x more total gene body dark bases (either type) for each technology, compared to CHM13, except ONT which was almost even (Illumina100: 1.97x; Illumina250: 2.25x; PacBio: 3.27x; ONT: 1.15x; **Fig. 2ci**). PacBio also had between 2.4x (CHM13) and 6.7x (HG38 Alt) more total gene body dark bases than ONT across all four reference genomes. We additionally observed that HG38 with alternate contigs has greater total gene-body dark-by-MAPQ bases in PacBio data (20.4 Mb) compared to HG38 without alternate contigs (2.8 Mb) and CHM13 (6.1 Mb), respectively, demonstrating the alignment challenges that the alternate contigs cause (**Fig. 2c**). One reason this may affect PacBio more than ONT is that PacBio has shorter read lengths than ONT (PacBio median read length, averaged across samples: 15,577bp; standard deviation: 1,495; ONT median read length, averaged across samples: 24,798bp; standard deviation: 9,769; **Supplementary Fig. S1e,f**). This trend also held when subsetting the genes to only protein coding genes (**Fig. 2d**), showing that CHM13 provides both a complete genome and avoids over representing gene regions. Significant work is still needed for CHM13 to be ready for widespread use, however. Specifically, the background work already done on previous genome assemblies, including allele frequencies and gene annotations, is not easily converted to a new reference genome (accurately).

After comparing the total gene body dark bases, we assessed the number of genes containing any dark region with at least 20 contiguous nucleotides. Exactly 8,077 genes contain dark regions (of either type) in short-read data aligned to CHM13, whereas ONT only had 286 (Illumina100: 8,077; Illumina250: 6,377; PacBio: 990; ONT: 286; **Fig. 3a**). Of the 8,077 genes containing dark regions, 6,059 contained dark-by-MAPQ (Illumina100: 6,059; Illumina250: 4,255; PacBio: 824; ONT: 180; **Fig. 3b**) and 3,709 genes contained dark-by-depth regions in CHM13 (Illumina100: 3,709; Illumina250: 3,372; PacBio: 319; ONT: 107; **Fig. 3c**). For clarity, the sum of dark-by-depth and dark-by-MAPQ gene counts add up to 9,768 for the 100-bp Illumina data (not 8,077) because 1,354 genes contain both dark-by-depth and dark-by-MAPQ regions within the gene body (Illumina100: 1,354; Illumina250: 907; PacBio: 57; ONT: 4; **Fig. 3d**).

**Figure 3.**
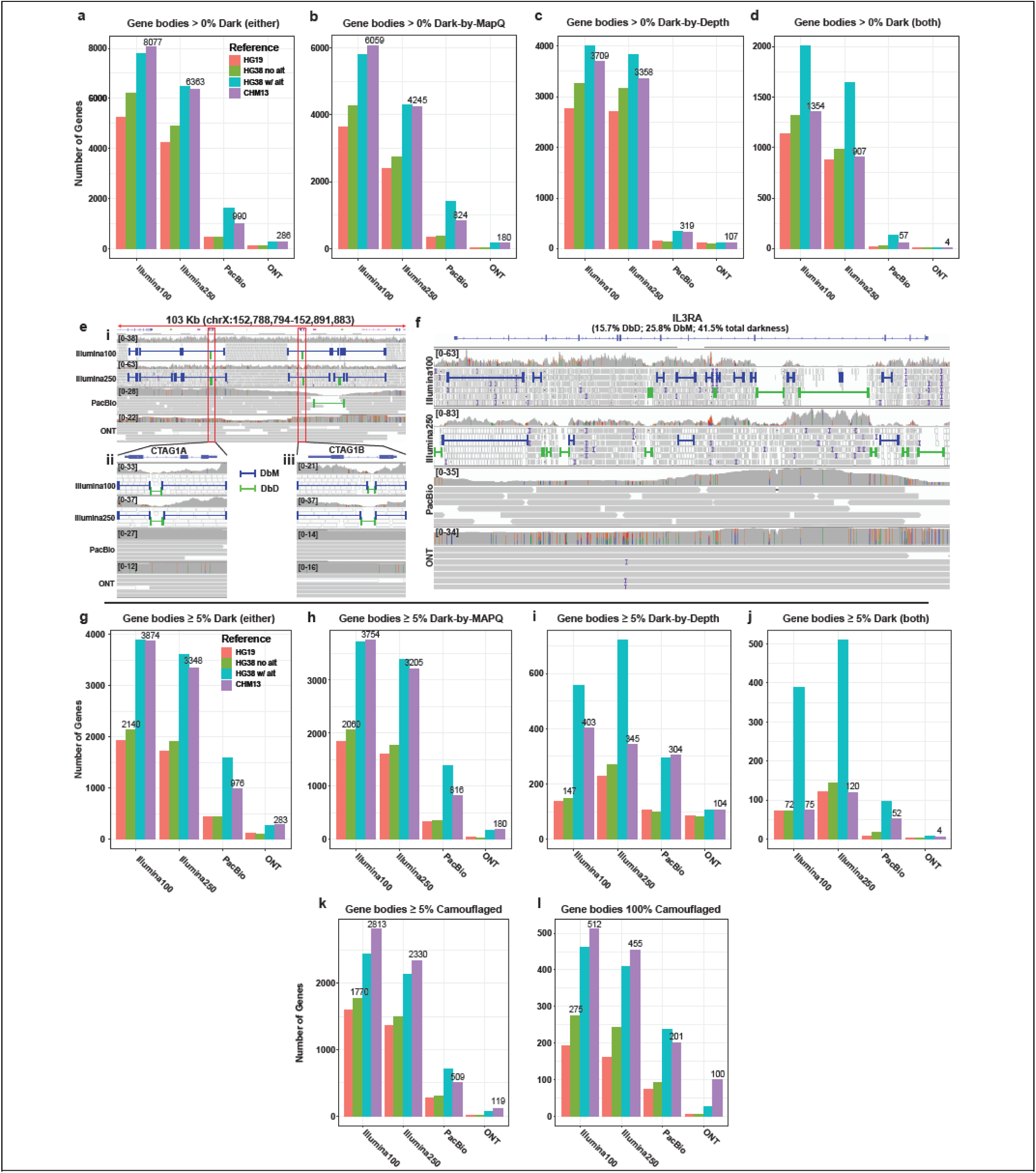
The T2T-CHM13 genome contained more dark and camouflaged genes than HG38; ONT outperformed other platforms. (**a**) 8,077 gene bodies contained dark regions in CHM13 for Illumina100 while ONT samples only had 286 dark gene bodies. (**b**) 6,059 gene bodies contained dark-by-MAPQ regions in CHM13 for Illumina100 while ONT samples only had 180 dark-by-MAPQ gene bodies. (**c**) 3,709 gene bodies contained dark-by-depth regions in CHM13 for Illumina100 while ONT samples only had 107 dark-by-MAPQ gene bodies. (**d**) 1,354 gene bodies contained both dark-by-depth *and* dark-by-MAPQ regions in CHM13 for Illumina100 while ONT samples only had 4 gene bodies containing *both* dark-by-depth and dark-by-MAPQ regions. (**e**) *CTAG1A/B* contained both dark-by-depth and dark-by-MAPQ within a larger dark region. (**f**) *IL3RA* is 41.5% dark (in Illumina100) made up of both types of dark regions. (**g**) CHM13 had almost double the number of at least 5% dark genes compared to HG38 without alternate contigs. (**h**) While CHM13 had far more genes with at least 5% dark-by-MAPQ, ONT resolved 95%. (**i**) Genes that were at least 5% dark-by-depth were an order of magnitude less of an issue (see Y-axis) than dark-by-MAPQ, but ONT still performed the best across the board. (**j**) HG38 with alternate contigs had more gene bodies with at least 5% dark by both types. (**k**) CHM13 generally had the most genes with at least 5% camouflaged, the vast majority of which were resolved with long reads. (**l**) CHM13 generally had the most 100% camouflaged genes.

Examples of genes that suffer from both forms of dark regions include *CTAG1A*, *CTAG1B*, and *IL3RA*. *CTAG1A* and *CTAG1B* (both originally known as *NY-ESO-1* before the duplication was discovered) are located in a duplicated region of chromosome X causing two approximately 35kb dark-by-MAPQ regions (**Fig. 3e.i**) and are primarily expressed in specific cell types within the testes and in several types of cancer (e.g. breast cancer, leukemia, etc.) making it a potential therapeutic target [19–23]. Problematically, *CTAG1A* and *CTAG1B* are 97.9% dark (dark-by-depth: 23.5%, dark-by-MAPQ: 74.4%; **Fig. 3e.ii**) and 99.6% dark (dark-by-depth: 26.9%, dark-by-MAPQ: 72.8%; **Fig. 3e.iii**) in Illumina100 sequencing aligned to CHM13, respectively. *CTAG1A* contains a dark-by-depth region in the first intron for both Illumina100 and Illumina250. *CTAG1B* also contains a dark-by-depth region in the intron which breaks up the dark-by-MAPQ regions that span the rest of the gene in Illumina100 and Illumina250 samples. PacBio sequencing resolved most of this large, duplicated region on chromosome X, while ONT resolved the entire region (**Fig. 3e.i**). While only ONT was able to fully resolve the entire region, PacBio had better coverage in *CTAG1A* and always has better single molecule per-base accuracy. *CTAG1A* and *CTAG1B* are perfect examples of genes and regions that prevent researchers from performing a truly complete genomic analysis, because short reads cannot be accurately disentangled across the region. Long-read data properly illuminate these dark regions.

*IL3RA* (a.k.a. *CD123*) is a subunit of a cytokine receptor with two copies, with one located on the X and Y chromosomes, each. It is a top therapeutic target for acute myeloid leukemia and blastic plasmacytoid dendritic cell neoplasm because it is known to have higher expression in leukemic cells [24–27]. *IL3RA* has a fascinating and complicated short-read alignment structure, oscillating from non-dark to dark-by-MAPQ or dark-by-depth (**Fig. 3f**). In total, it is 41.5% dark with a combined 25.8% of the gene being dark-by-MAPQ and 15.7% of the gene being dark-by-depth. The sporadic dark regions across *IL3RA* using short-read data prevent a complete analysis of this gene, which is believed to be key to treating leukemic cells. PacBio has decreased coverage at the ends of *IL3RA* but better single molecule per-base accuracy, whereas ONT had more consistent, deep coverage across the gene. By leveraging long-read data, a complete analysis of this gene is possible and could provide additional clues for treating disease.

Looking at gene bodies that are at least 5% dark (of either type), we observed 3,874 when aligning Illumina100 reads to CHM13, which is nearly double the number of gene bodies that were 5% dark when aligned to HG38 without alternate contigs (2140), and nearly four and fourteen times more than PacBio (976) and ONT (283) when aligned to CHM13, respectively (**Fig. 3g**). Exactly 3,754 (96.9%) of the dark genes contained dark-by-MAPQ regions in CHM13 (**Fig. 3h**). ONT resolved 95.2% 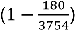 of the Illumina100 dark-by-MAPQ genes, while Illumina250 only resolves 14.4% and PacBio resolves 78.3% in CHM13 (**Fig. 3h**). When stratifying by dark-by-depth (**Fig. 3i**) or gene bodies that contain both types (**Fig. 3j**), HG38 with alternate contigs stands out with the greatest number of gene bodies that are at least 5% dark in short-read data, as expected; we expected HG38 with alternate contigs to have more dark gene regions because the alternate contigs introduce additional gene copies. The trends remain similar when comparing genes that are 100% camouflaged (**Fig. S2a-d**). Long-read sequencing technologies outperform the short-reads in all categories, and ONT outperforms PacBio.

To understand the difference between references, we assessed the number of camouflaged genes that are at least 5% camouflaged. We also assessed only those that are 100% camouflaged, specifically. For reference, camouflaged genes are dark-by-MAPQ regions that fall within a gene body and have at least 98% similarity to another genomic region (i.e., a subset of dark-by-MAPQ). Exactly 2,813 genes have more than 5% of the gene body camouflaged in Illumina100 data when aligned to CHM13 (Illumina100: 2,813; Illumina250: 2,330; PacBio: 509; ONT: 119; **Fig. 3k**) and 512 gene bodies were 100% camouflaged (Illumina100: 512; Illumina250: 455; PacBio: 201; ONT: 100; **Fig. 3l**). The 512 gene bodies that were 100% camouflaged with Illumina100 reads aligned to CHM13 is nearly double than when aligned to HG38 without alternate contigs (275). These gene bodies were predominantly classified pseudogene, protein coding, and lincRNA biotypes in those that are greater than 5% and 100% camouflaged (**Fig. S2e,f**).

ONT can further elucidate the role of large variants in dark reads where even PacBio struggles. An example is the intron of *TMEM88B,* which contains a dark-by-depth region flanked by dark-by-MAPQ regions in the Illumina100 data (49.5% dark-by-depth; **Fig. S2g**). The long-read data decreases the size of the dark region, but only ONT unequivocally resolved the region, showing that this subject likely has a heterozygous deletion within the *TMEM88B* intron. Little is known about TMEM88B, though it appears to be a negative regulator of WNT signaling, particularly in the heart [28, 29]. Thus, the effect of this intragenic deletion is undetermined, but worthy of future research. Many more ONT reads properly map to the general region (46; **Fig. S2g**) compared to PacBio (10) despite similar overall sequencing depth, and many more ONT reads spanned the entire region. The heterozygous deletion is indicated by approximately half of the ONT reads mapping through the entire region containing a deletion, while the other half map through the region without a deletion. Thus, long reads suggest that the short-read alignment challenges in the *TMEM88B* intron are likely due to a large intronic deletion. *TMEM88B* highlights the utility of long-read data for identifying relatively large deletions and therefore further the quest for a truly complete analysis by shedding light on the dark regions of the genome.

### Dark regions from short reads obscure short-read DNA epigenetic assay results

After quantifying genome-wide dark and camouflaged regions, we anticipated these dark regions would likely be consistent across the spectrum of genomic DNA assays, including those related to epigenetics. Specifically, we looked at bisulfite sequencing, chromatin immunoprecipitation (ChIP) sequencing, and high-throughput chromosome conformation capture (HiC) sequencing. Whole-genome bisulfite sequencing is a common assay to assess the CpG methylation status genome wide by converting unmethylated cytosines to uracils (**Fig. 4a**). ChIP-Seq, on the other hand, is an assay designed to identify DNA binding locations for proteins to a specific place in the genome (**Fig. 4b**). HiC data is an epigenetic assay that identifies regions of the genome that physically interact and is used to identify boundary regions within which an enhancer or other distal regulatory element could interact with a given promoter called Topologically Associated Domains (TADs; **Fig. 4c**) [30]. All three assays historically rely on short-read sequencing and are thus likely susceptible to issues related to dark and camouflaged regions. To test the effect dark regions have on epigenetic assay results, we identified dark bases and camouflaged genes in each assay and found that epigenetic assays are equally susceptible to challenges related to dark regions.

**Figure 4:**
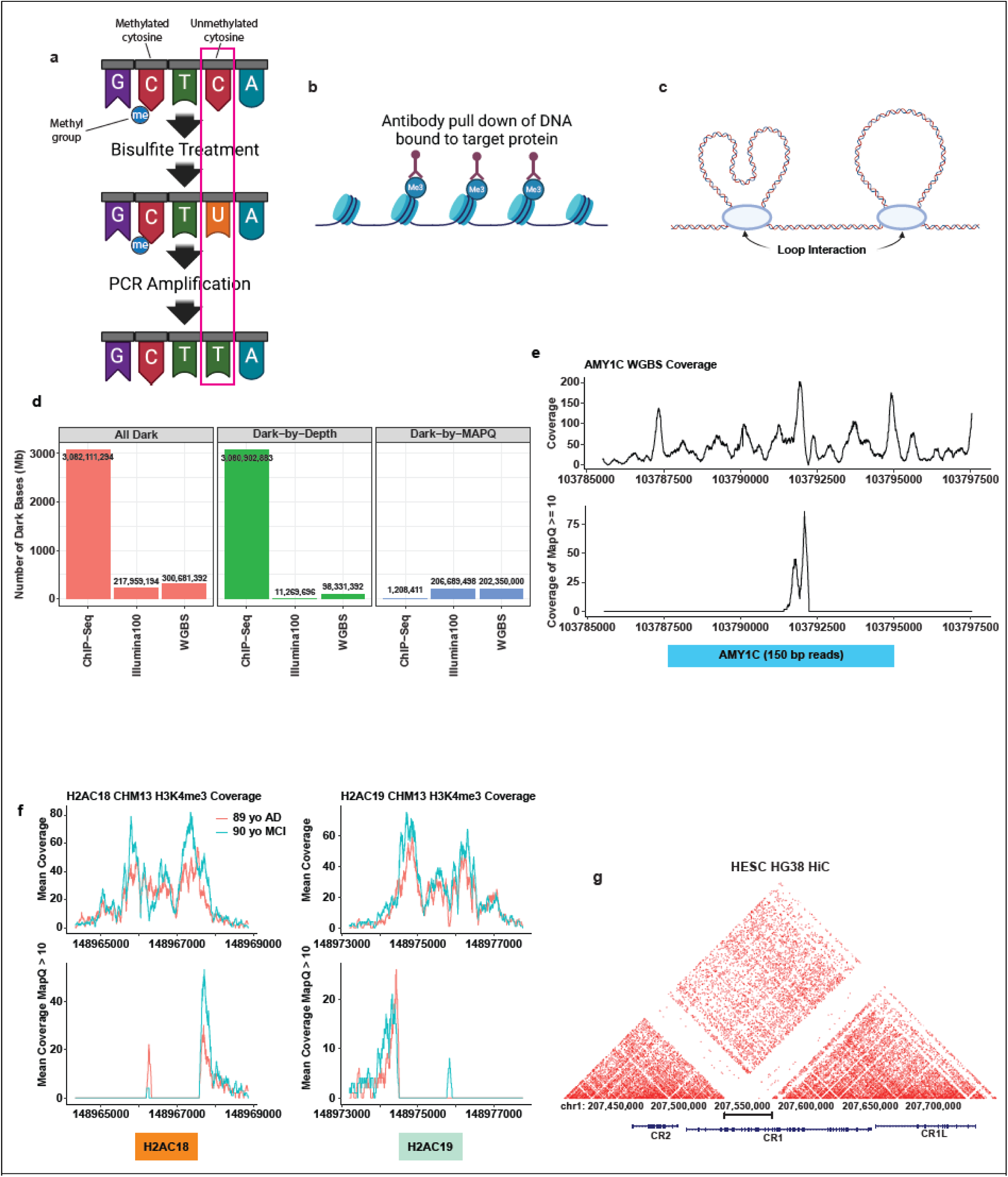
Dark regions obscure epigenetic assay results. **(a)** Whole genome bisulfite sequencing is a common genome-wide short-read sequencing assay that identifies CpG methylation sites by converting unmethylated cytosines to uracils. **(b)** ChIP-Seq is an assay that pulls down DNA bound to a target protein. **(c)** HiC data identifies the location of chromatin loops in nuclear DNA. **(d)** Compared to 100 bp Illumina data, the whole-genome bisulfite sequencing samples have more dark-by-depth and less dark-by-MAPQ. Most of the ChIP data, as expected, are dark-by-depth; however, 1.2 Mb (3.4%) are dark-by-MAPQ. **(e)** Due to close paralogs few, if any, CpG sites in *AMY1C* can be quantified using short-read sequencing because *AMY1C* is dark-by-MAPQ and camouflaged by *AMY1A* and *AMY1B*. **(f)** *H2AC18/19* camouflage each other and, as a result, the promoter associated histone mark H3K4me3 peaks are obscured and left unanalyzed. **(g)** The *CR1* tandem domain repeats completely obscure DNA looping data in HiC data.

We compared the difference in total dark bases, dark-by-depth bases, and dark-by-MAPQ bases with whole-genome bisulfite sequencing, ChIP-seq, and the previously analyzed Illumina100 dark data. ChIP-seq assays will only include DNA bound to the queried proteins (e.g., H3K4me3 histone marks), thus we expect most of the genome to be dark-by-depth, by nature of the assay. We leveraged frontal cortex (BA9) H3K4me3 promoter associated histone mark data [21, 31] for two individuals we obtained from the Encode Project. H3K4me3 dysfunction is linked to several neurological conditions including Alzheimer’s disease [32], Autism [33, 34], and Schizophrenia [35]. As expected, we found the ChIP data had far more dark bases than both Illumina100 and whole-genome bisulfite sequencing. For those regions where coverage is expected, however, high-quality alignments are essential. Of the approximately 35 megabases (35,180,776 bp) where ChIP-seq sequencing data were obtained, 1.2 megabases (3.4%) of ChIP data were dark-by-MAPQ (**Fig. 4d**). Thus, even those regions where sequencing data are expected, short-read ChIP-seq results leave a large proportion inaccessible because they are dark-by-MAPQ.

To assess the effect of dark regions on whole-genome bisulfite sequencing data, we leveraged publicly available data from the Encode Project from male adrenal gland tissue, where the sequencing read length was 150 bp. Even knowing that bisulfite treatment is extremely harsh on DNA, we observed a surprisingly large number of dark-by-depth bases, resulting in an 8.7x increase (98,331,392 bp) compared to Illumina100 whole-genome sequencing (**Fig. 4d**). Such a large number of bases without sufficient coverage precludes the ability to assess DNA methylation. While the increase in dark-by-depth is concerning, it is not necessarily surprising because harsh bisulfite treatment shears the DNA, and biases are well known [36–38]. Finally, about 202 megabases (202,350,000 bp) of the whole-genome bisulfite sequencing data are dark-by-MAPQ. We conclude that short-read based epigenetic assays are dramatically affected by dark regions, predominantly dark-by-depth bases (**Fig. 4d**).

Here, we highlight an example camouflaged gene for each of the epigenetic assays. First, we investigated the whole-genome bisulfite sequencing camouflaged genes. We plotted the per base genomic total coverage and high-quality coverage (MAPQ ≥ 10) for the *AMY1C* gene, which is 49.6% camouflaged in Illumina100 data and whose copy number relates to glucose absorption [39, 40] (**Fig. 4e**). The promoter proximal region of *AMY1C* has sufficient total depth to assess CpG methylation (**Fig. 4e** top); however, except for one small region of the gene, the region is dark-by-MAPQ (**Fig. 4e** bottom), and therefore is unrepresented in the processed data. Using short-read bisulfite sequencing data, it is not possible to assess the epigenetic regulation of the *AMY1C* gene by DNA methylation using standard analysis pipelines. An awareness of dark-by-MAPQ regions in genes is prudent when assessing CpG methylation. In our original paper [1], we developed a method to rescue and analyze reads from dark-by-MAPQ regions, but we maintain that this approach is simply a stopgap and long reads are the ultimate solution—especially since long-read technologies can directly measure genome-wide DNA methylation during standard sequencing without special DNA preparation.

Secondly, we highlighted several genes in ChIP-Seq data that are camouflaged by each other, including the histone genes *H2AC18* and *H2AC19* (100% and 99.8% camouflaged respectively in Illumina100; **Fig. 4f**), and the heat shock proteins *HSPA1A* and *HSPA1B* (41.4% and 39.5% camouflaged respectively in Illumina100; **Fig. S3a**). For both sets of genes, we saw large peaks in the total coverage in the promoter proximal region of these genes; however, the peaks in the promoter proximal region are truncated or completely obscured by large dark-by-MAPQ regions. While little is known about the differential expression of *H2AC18* and *H2AC19*, they are differentially expressed upon exposure to a Human Papillomavirus oncoprotein [41] therefore an understanding of how these genes are epigenetically regulated has potential medical importance. *HSPA1A* and *HSPA1B* are two subunits of the HSP70 complex with potential therapeutic relevance to ALS [42–44]. When camouflaged regions obscure epigenetic results for potentially medically relevant genes such as *H2AC18/19* and *HSPA1A/B*, a complete understanding of gene regulation and a comprehensive analysis is inhibited.

Finally, HiC data is also plagued by gaps due to dark regions because it relies on short-read sequencing. For example, the Alzheimer’s disease-associated gene, *CR1*, has a tandem domain duplication that camouflages a large segment of the gene [1]. This region is almost completely dark in HG38 HiC data (**Fig. 4g**). Meng *et al.* identified an overlap of Alzheimer’s disease associated GWAS SNPs with HiC interactions and eQTLs [45], but their results would likely overlook any association within *CR1* because the region is camouflaged. The use of TADs through the HiC assay in Alzheimer’s disease would leave the analysis blind to any physical interactions or GWAS SNPs with this region of *CR1* and it would increase the complexity of TAD boundary identification.

Ultimately, a complete analysis requires the integration of multiple assays to fully understand genomic processes. The loss of valuable information from these epigenetic assays because of short-read sequencing inhibits a comprehensive genomic analysis. The inability to quantify DNA methylation, histone modifications, and genomic looping causes a gap in our understanding of gene regulation and overall genomic structure of these regions. In addition to largely resolving camouflaged regions of the genome through long-read sequencing, PacBio and ONT have the added benefit of being able to identify DNA methylation simultaneously.

### CHM13 results in decreased dark bases in CR1 resulting in better identification of CR1 major allele

Insertions and deletions can modify copy number which can influence disease development for a range of diseases, including Alzheimer’s disease (*APP* duplication) [46], Parkinson’s disease (*SNCA* duplication) [47, 48], and various heart diseases [49, 50]. Previous work has even suggested that copy number of the *CR1* C3B/C4B binding domain is associated with Alzheimer’s disease risk [51]. The *CR1* C3B/C4B binding domain is known to be variable in the population [52]. Specifically, *CR1* is known to have at least four primary haplotypes known as *CR1*-A, *CR1*-B, *CR1*-C, and *CR1*-D [51], where all four haplotypes are believed to originate from varying combinations of three highly similar low-copy repeats (LCR) known as LCR1, LCR1’, and LCR2 that make up the C3B/C4B binding domain and plays an important role in the complement cascade.

At the protein level, all three LCRs are identical except for a single amino acid difference in LCR1 (Ala405Thr). Thus, treating these three repeats as the same fundamental domain, HG38 contains three total copies of the binding domain [1], whereas CHM13 contains only two [52] (**Fig. 5a,b**). Yang *et al.* recently demonstrated that the *CR1* allele represented in CHM13 (two copies) is the major allele found in 79 of their 94 samples (84%), while HG38 contains the minor allele (three copies) [52]. The ramifications of the reference genome used when analyzing sequencing data for this region are important. Because the binding domain is repeated, it is also dark-by-MAPQ (and camouflaged) in short-read sequencing data, as we demonstrated previously [1]. When analyzing Illumina100 data for this region using HG38, *CR1* is 22.7% camouflaged, whereas it is only 3.1% camouflaged when using CHM13. This dramatic difference is not only because one of the copies is missing, but because the *CR1*-A allele contains the two most distinct LCRs (LCR1 and LCR2) [53–55]. Analyzing short-read data in CHM13 makes it possible to more easily identify variants that would be obscured by HG38 but will also result in more false variants being reported for individuals carrying the *CR1*-B allele because of true differences between the three LCR domains at the nucleotide level. This shows that the reference genome you use matters when assessing regions with variable copy numbers in the population in short read data. Notably, both PacBio and ONT maintain coverage through this region regardless of the reference genome used.

**Figure 5:**
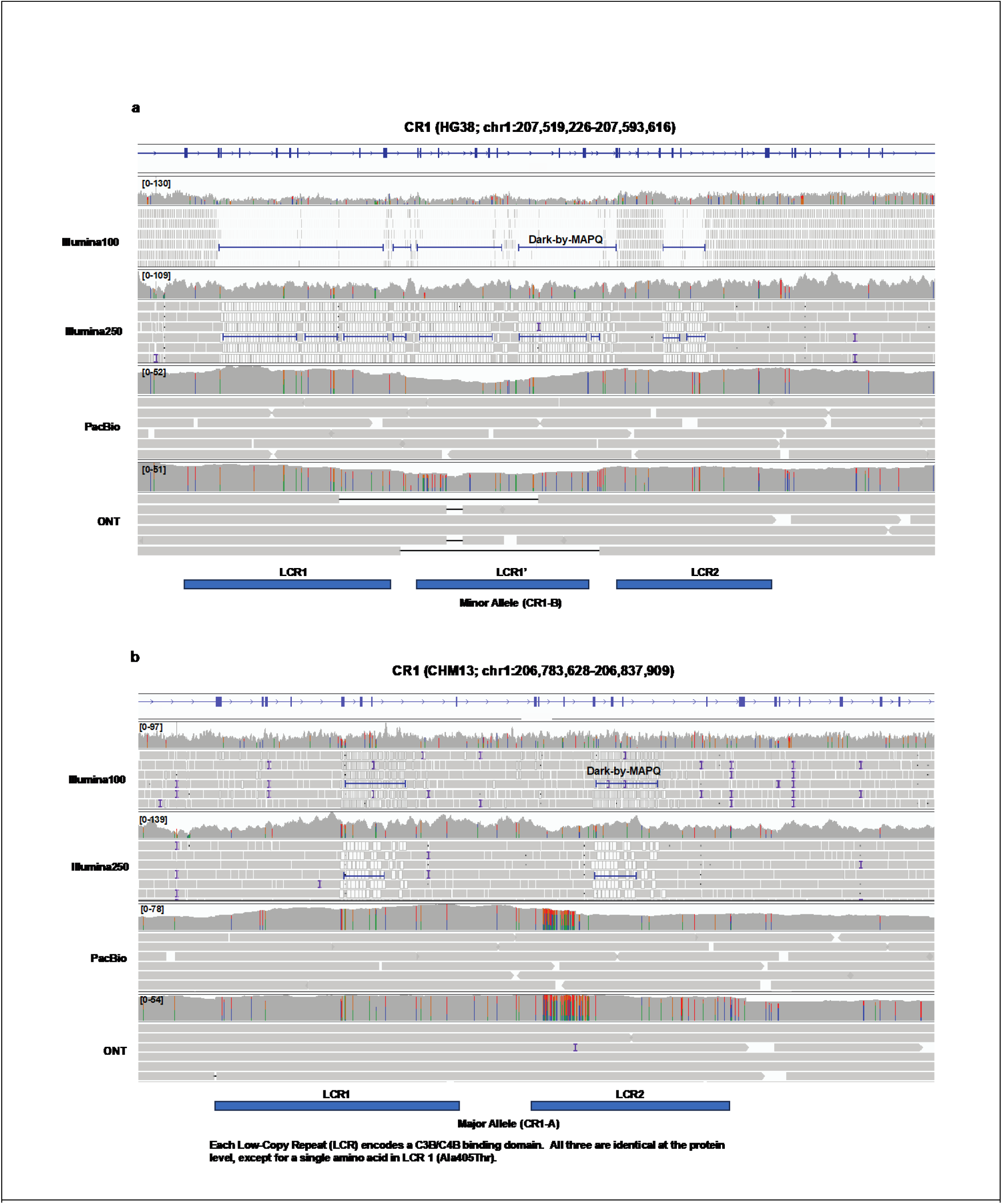
*CR1* tandem C3B/C4B domain repeat number differs between HG38 and CHM13 with important ramifications for short-read alignments while long reads perform well on both. *CR1* haplotypes are made up of varying numbers of low-complexity repeat regions (LCRs). **(a)** HG38 contains the *CR1*-B haplotype with three copies of the C3B/C4B binding site domain representing the minor allele where we see widespread dark-by-MAPQ (camouflaged) regions with short-read data. *CR1* is 22.49% camouflaged in HG38 (including introns). Both PacBio and ONT resolve this region well. **(b)** CHM13 contains the *CR1*-A haplotype with two copies of the C3B/C4B binding domain. The amount of dark-by-MAPQ (camouflaged) bases is dramatically fewer, where *CR1* is 3.07% camouflaged in CHM13.

### Centromeric satellite variability on chromosome 10 drives poor alignments for both short- and long-read data in CHM13

We were curious about the nature of genomic regions where short-read data exhibit both dark-by-MAPQ and dark-by-depth behavior. We identified a large region in CHM13 near the chromosome 10 centromere that exhibited this behavior (**Fig. 6a**). We found that the region is highly repetitive with multiple repeat types, including human satellite II (HSATII) and Beta satellite sequences (**Fig. 6a**). Most of this region is dark-by-MAPQ, which is easily explained because of ambiguous alignments from sequence duplication. We anticipate that the dark-by-depth regions could only occur for two reasons: (1) there simply are no reads from that region for the individual (e.g., genuine genomic deletions, sequencing artifacts, etc.); or (2) because of an alignment issue.

**Figure 6:**
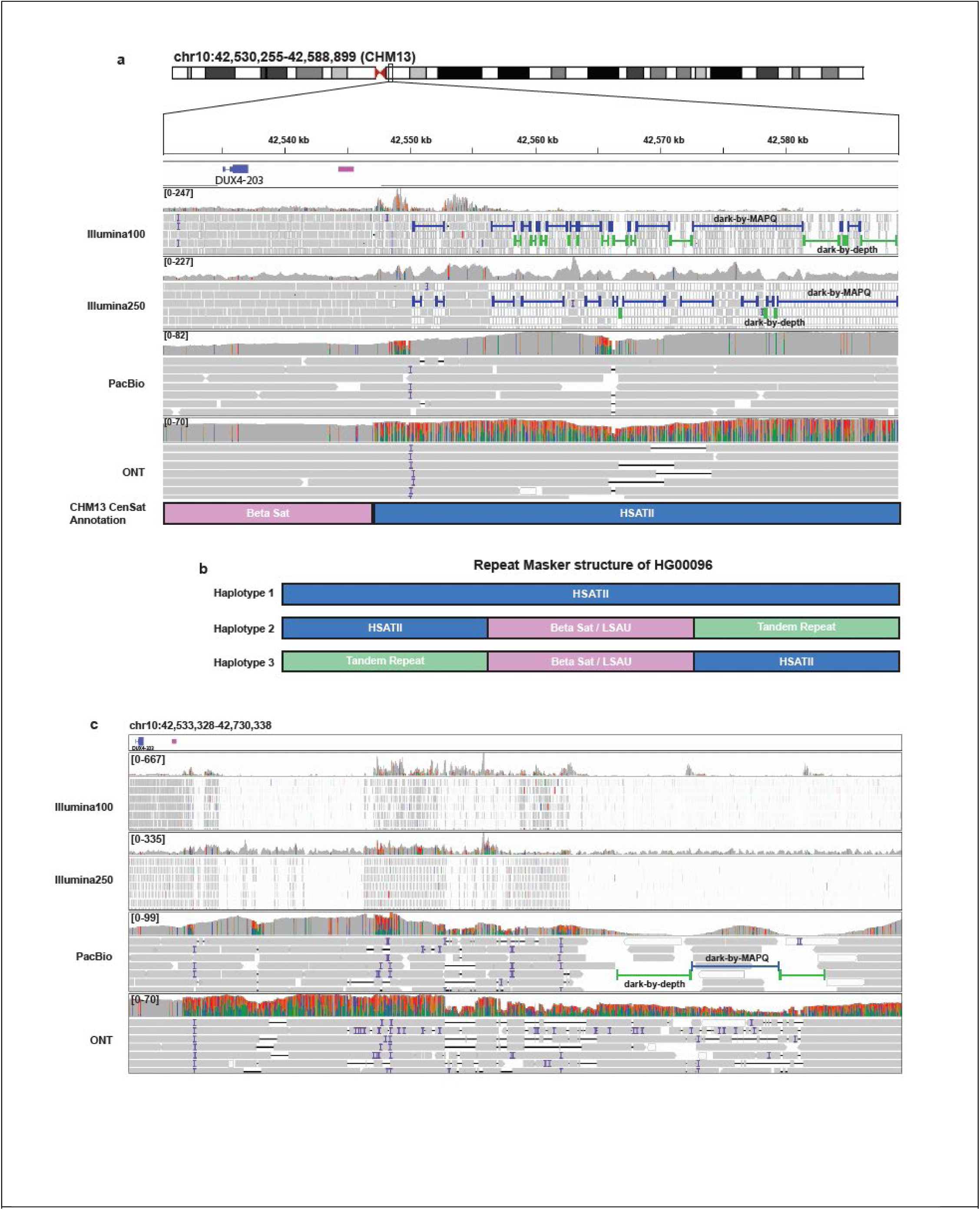
Centromeric/pericentromeric satellites cause problems in alignments. **(a)** The pericentromeric region exhibits major alignment challenges with varying degrees of dark-by-MAPQ and dark-by-depth regions. **(b)** Using Repeat Masker, we identified multiple types of repetitive elements rather than just the annotated HSATII. **(c)** When zooming out we see that ONT has more deletions and insertions, combined with a large section with nearly complete sequence variability that still aligns (indicated by high mismatch rate in histogram above the ONT reads). On the other hand, PacBio has large dark-by-depth regions.

In addition to the co-occurring dark-by-MAPQ and dark-by-depth regions within short-read data, we noticed that the long-read alignments were also problematic. Part of what makes this example particularly striking is the clear contrast between the PacBio and ONT alignments. For PacBio, a large portion of this region is well represented with quality alignments but reaches a sudden dark-by-depth region that is not observed in ONT data (**Fig. 6a**). On the other hand, while ONT maintains better coverage throughout the region, there is a sudden drop in alignment quality not observed in the PacBio data that coincides with a known boundary between a Beta satellite and HSATII (**Fig. 6a**). The drop in ONT alignment quality is shown by the sudden increase in mismatches in the histogram above the ONT reads (**Fig. 6a**).

To better understand this phenomenon, we collected all ONT reads that aligned to this region (chr10:42,530,255-42,588,899) for this sample (HG00096) and analyzed them individually using RepeatMasker after converting all the reads to the same forward direction. Based on the RepeatMasker results, we were surprised to find three different haplotypes in this region: (1) the first consisted solely of HSATII sequence, matching the reference genome (57/80 reads; 71.25%); (2) the second contained the annotated HSATII followed by a combination of beta satellites, LSAU, and composite beta/LSAU repeats, and tandem repeats (HSATII/BSat/LSAU/TR; 11/80; 13.75%); and (3) the third was an inverted version of the second (12/80; 15%; **Fig. 6b**). The composite beta/LSAU repeats were identified during the T2T studies [16], and this region appears to include families 1,4, and 10. Observing three haplotypes is entirely unexpected and suggests there may be mosaicism occurring in this region. This finding deserves to be followed up in future studies. Notably, Altemose *et al.* recently reported that satellites make up 6.2% of the CHM13 reference genome, and specifically discussed the chromosome 10 centromere as being highly variable because of structural variation [9]. We also noticed that one of the two known copies of the *DUX4* gene is located in this region. A repeat retraction in both the chromosome 4 [56] and 10 [57] copies are known to cause facioscapulohumeral muscular dystrophy (FSHD). Looking at alignments within a broader region, we see a decrease in general coverage over this centromeric region and an increase in dark regions and insertions and deletions in long-read sequencing pointing to incomplete alignments likely due to the satellite rearrangements (**Fig. 6c**). In all, these results suggest that centromeric satellite variability drives alignment challenges for both short- and long-read data—perhaps because using a static reference genome imposes the reference genome’s structure on the individuals.

### Supplementary alignments resolve some dark regions in long-read data

During our analyses, we identified dark regions in long-read data aligned to CHM13 overlapping known structural variants that were properly resolved with short-read data. One example is on the q-arm of chromosome 8 (chr8:112,191,464-112,192,351), corresponding to a documented inversion in CHM13 compared to HG38 with the Cactus [58] alignment annotations found on the UCSC Genome Browser [59] (**Fig. 7a**). We identified this region because of a stark, consistent dark-by-depth region in all long-read samples aligned to CHM13 (using only primary alignments). This region of 887 bases is completely dark and ends abruptly (**Fig. 7b**). In this situation, the reads were long enough to span such a short region (887 bases), thus it is unlikely that the long-read data itself was the problem. Short-read sequencing data clearly identified the inversion where paired-end reads are oriented in the same direction (highlighted reads; **Fig. 7b**) with a break in the coverage at the breakpoints of the inverted region. The reads for all ONT and PacBio samples were soft clipped to exactly match the same boundaries. Thousands of bases were soft-clipped per read in PacBio, and tens of thousands of bases were soft-clipped per read in ONT. Notably, this gap was not present in HG38 alignments (**Fig. 7c**).

**Figure 7:**
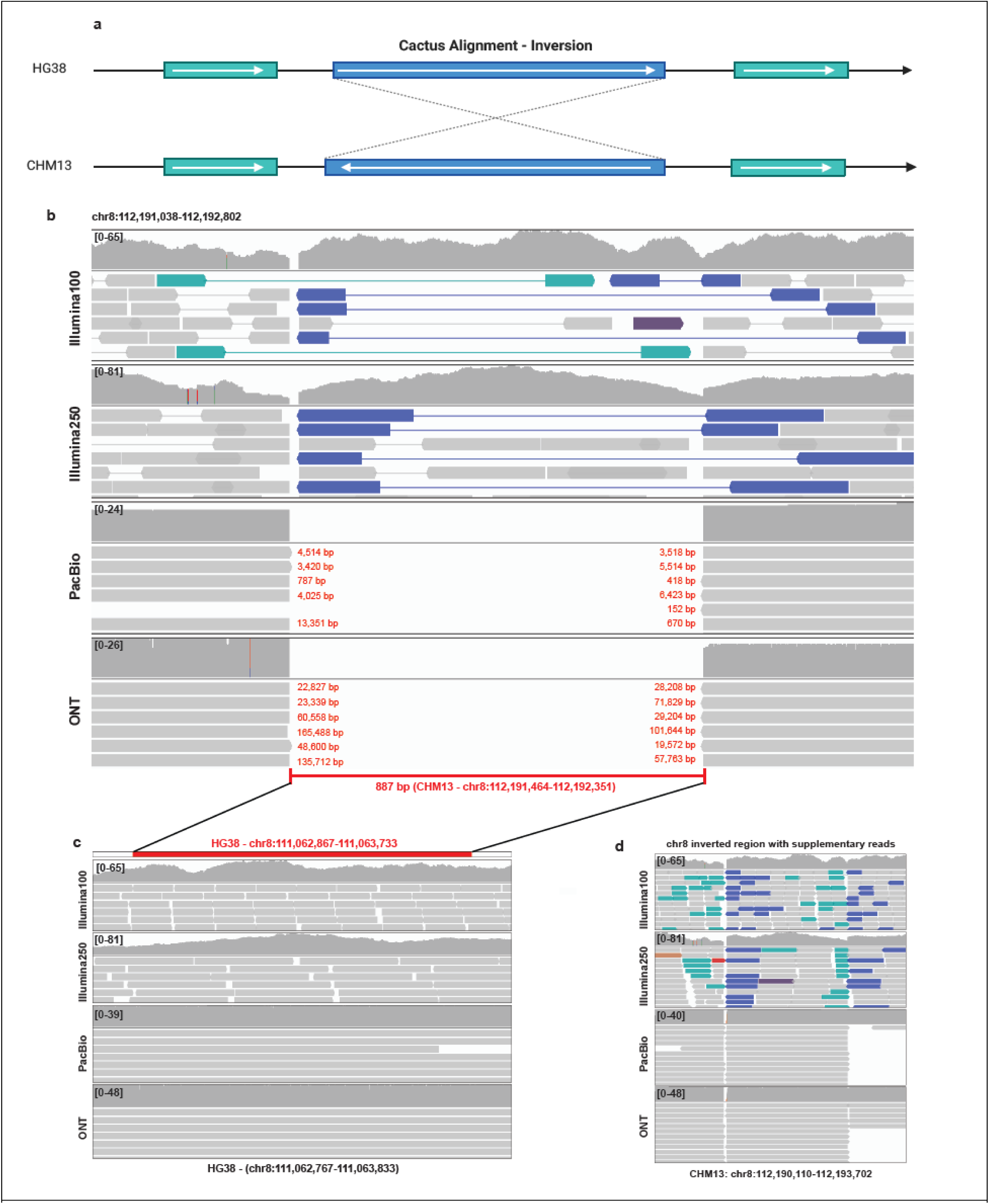
Inversions cause decreased coverage in long read alignments. **(a)** Chromosome 8 contains a region of 887 bp that is consistently dark-by-depth in all PacBio and ONT samples where minimap2 soft-clipped the reads and Cactus alignments show an inversion between HG38 and CHM13. **(b)** In CHM13, short-read alignments clearly identify the inversion, while primary long-read alignments (removing supplementary alignments) soft-clip large segments of reads. **(c)** In contrast to the CHM13 alignments, this region is not inverted in HG38 and is completely covered in all alignments. **(d)** Supplementary reads resolve dark region caused by only considering primary reads in the inverted region.

In our original paper [1], for simplicity we excluded all secondary and supplementary reads when assessing dark regions, therefore including only primary reads. In these analyses, we found that the break in alignments stems from our exclusion of supplementary alignments. Minimap2 uses supplementary alignments to span the breaks in the case of inversions. To test this, we re-analyzed the samples including the supplementary alignments and, in this case, including supplementary reads resolved this dark-by-depth region (**Fig. 7d**). To assess the overall effect of using supplementary reads, we determined to compare the differences between the primary only results and the primary and supplementary results.

When including supplementary alignments, we found an expected increase in dark-by-MAPQ bases in short-read sequencing data. **(Fig. S4a-f**; **Table 2**; **Fig. 8a)**. The number of dark-by-MAPQ bases across the genome only slightly increased in HG38 with and without alternates for every sequencing platform except ONT which exhibited a decrease of almost two megabases when including supplementary reads in the analysis (1,685,814 bases in HG38 no alternate contigs; 1,695,547 bases in HG38 with alternate contigs). In ONT specifically, supplementary alignments consistently resulted in fewer dark-by-MAPQ bases across all references, though surprisingly, PacBio had a significant reduction in CHM13, compared to ONT, with 4 and 1 Mb decreases, respectively (**Fig. 8a**; **Table 2**). Contrastingly, Illumina100 had almost a 2 Mb (1,911,880 base) increase in dark-by-MAPQ bases in CHM13 (**Fig 8a**). Including supplementary reads consistently reduced dark-by-depth regions for all platforms, especially for ONT aligned to CHM13 which decreased by 15.5 Mb (**Fig. 8b**; **Table 2**).

**Figure 8:**
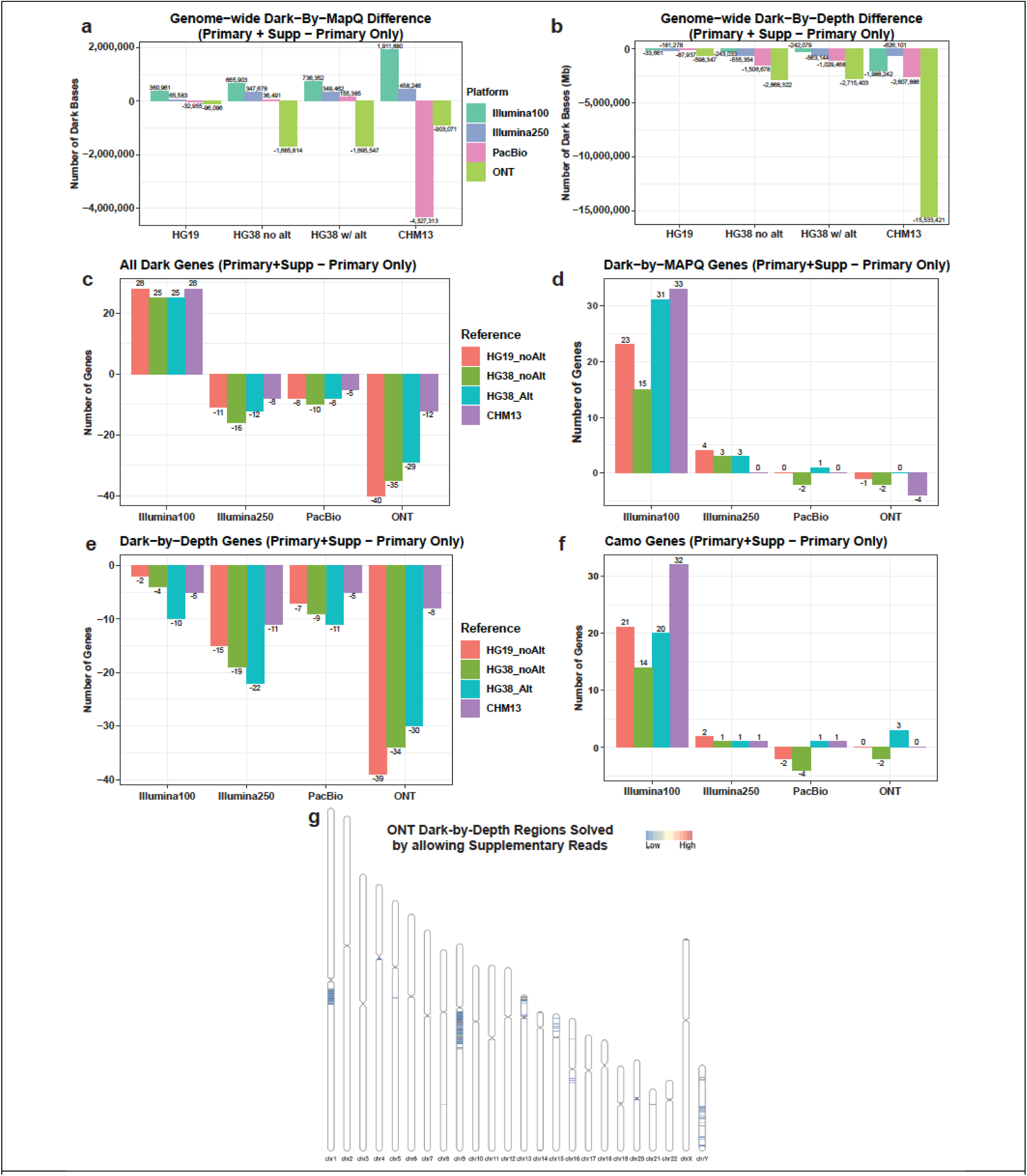
Adding supplementary reads to the process mainly effects long-read sequencing analyses. For each of the figures a-f a negative value means that adding in the supplementary reads resolved the dark regions, while a positive value means the dark regions are exacerbated. For figures c-f all genes were filtered based on having at least 5% of the gene being dark or camouflaged. **(a)** When comparing primary only alignments to primary with supplementary alignments included, ONT, in HG38 and CHM13, and PacBio, in CHM13, exhibit a large number of bases that are no longer dark-by-MAPQ. Illumina100 increased dark-by-MAPQ bases by almost 2 Mb. **(b)** ONT resolved the most dark-by-depth bases compared to all the other platforms across all references. **(c)** Including supplementary reads results in an increase in genes that are at least 5% dark in Illumina100 sequencing, and less for all other sequencing types. **(d)** Illumina100 has issues with increased numbers of at least 5% dark-by-MAPQ genes, while the other platforms change very little. **(e)** All platforms across all references have a decrease of genes that are at least 5% dark-by-depth. **(f)** Only Illumina100 saw an increase in genes that are at least 5% camouflaged with including supplementary reads. **(g)** We plotted the location of all of the ONT dark-by-depth regions resolved by including supplementary reads in the analysis.

**Table 2:**
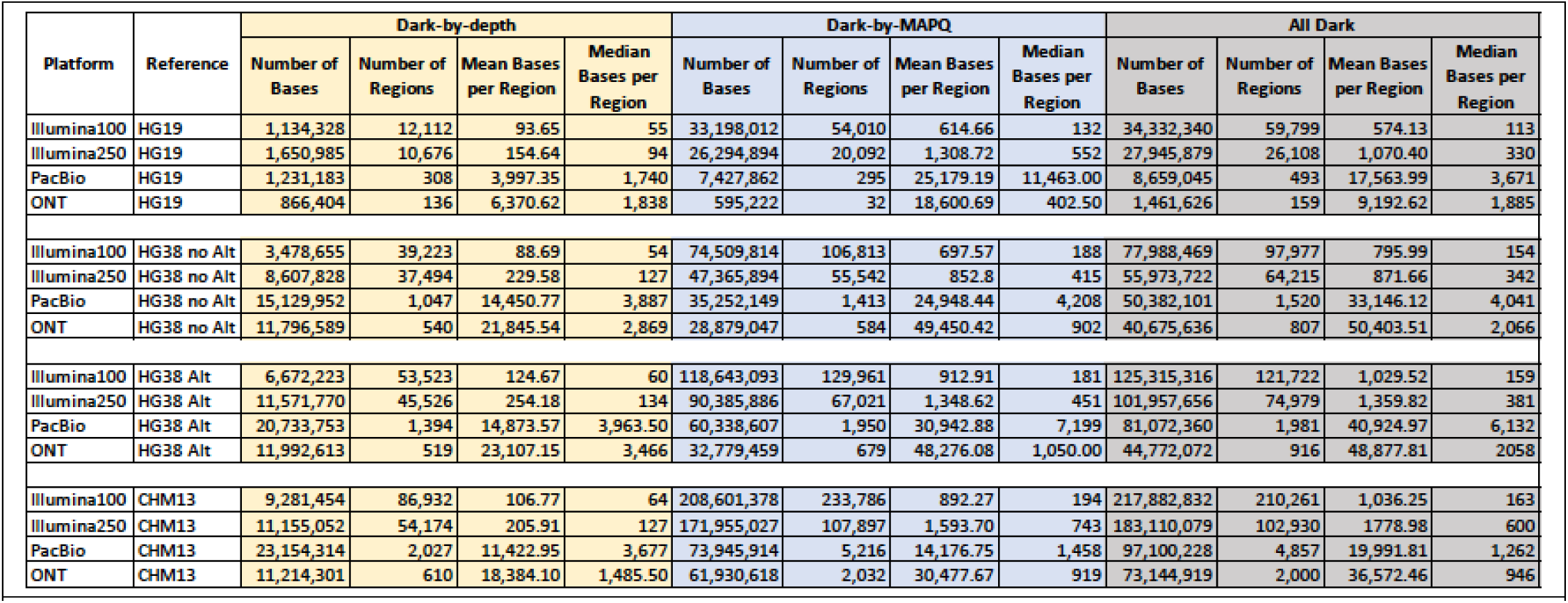
Dark bases and regions with supplementary read inclusion. When comparing this table to Table 1, dark regions and bases are mostly decreased across the board.

These differences in dark regions resulted in all sequencing platforms, except Illumina100, resolving dark regions within genes (**Fig. 8c**). We found that, across all platforms, dark-by-MAPQ had increased or no real change in the number of genes (**Fig. 8d**), while all platforms had a decrease in dark-by-depth genes when including supplementary reads (**Fig. 8e**). When looking at all dark genes, we observed an increase in Illumina100, while the others decreased. (**Fig. 8c**). All references across all the sequencing platforms exhibit a decrease in genes that are at least 5% dark-by-depth (**Fig. 8e**). Finally, only Illumina100 has an increase in genes that are at least 5% camouflaged when you add in supplementary reads (**Fig. 8f**). We plotted the location of the dark-by-depth regions in ONT sequencing that are resolved when including supplementary reads, revealing the biggest groupings are on the q-arms of chromosomes 1 and 9 (**Fig. 8g**).

These findings suggest that including supplementary reads results in marginal differences in short-read data while large, potentially significant differences, especially in structural variant identification, are found in long-read data, particularly ONT. This is likely because the increased read lengths often allow for spanning breakpoints that create chimeric reads.

## Discussion

Building off our previous work, in this study, we found that dark-by-MAPQ bases are the predominant dark regions across the spectrum of references—especially CHM13. Long-read platforms outperform short-read platforms in resolving dark regions, where ONT remains the most effective, including in gene bodies. Additionally, all short-read assay results are dramatically affected by dark regions, including epigenetic assays. Still, much work remains to achieve a comprehensive analysis, because even long-read technologies struggle to properly resolve highly variable regions when constrained to the structure of a static reference genome, particularly the highly variable centromeric satellites on chr10.

We also demonstrate the importance of including supplementary alignments with long-read data to properly resolve structural variants; the longer reads become the more necessary aligning reads chimerically becomes when aligning to a static reference genome. Finally, we also demonstrate that the reference genome can have important implications using *CR1* as a case example. From our analysis, we conclude that in many cases CHM13 and long-reads are likely the best choice, but in cases like addressing the minor allele of *CR1*, CHM13 may not be the best choice.

No single reference genome will be perfect. Based on certain statistics, HG19 may appear ideal since here we show that it has overall fewer dark regions than the other reference genomes—but it has fewer dark regions because it is incomplete. The more complete the human reference genome becomes, the more challenging it is to properly analyze and interpret an individual’s unique combination of DNA variants. These challenges are often due to genomic duplications and ambiguous alignments leading to dark and camouflaged regions, among other complications. Understanding what contributes to these challenges and how we can overcome them is imperative.

Our study further demonstrates the importance of long reads to achieve a comprehensive analysis but also shows that more work remains. The biggest limitation and future direction to this analysis is that we did not include the human pangenome project. The pangenome is the first reference genome that begins to account for population variability rather than attempting to represent all of humanity using a single, static reference genome. Therefore, the next logical step is to assess how dark regions impact the pangenome.

Ultimately, many challenges remain to realizing a comprehensive and personalized genomic analysis, including having a deeper understanding of the human genome itself, which is limited by imperfect sequencing and downstream analyses. Even a relatively basic comparison between the various reference genomes and respective sequencing technologies becomes challenging. We demonstrated a range of differences between the issues related to short- and long-read sequencing, but even important differences between long-read technologies and their tendencies. There remain significant gaps in our understanding of the human genome that limit our ability to study it. The problem becomes significantly harder when trying to account for genomic diversity across the population. Having the first truly complete human reference genome is an important step forward, but significant work remains.

## Methods

### Sample Selection

For our dark region analyses, we selected 10 samples for each type of data. We used the same short-read samples as we used in our 2019 analysis. For 100 bp read length Illumina data, we selected 10 male hispanic non-related samples from the Alzheimer’s Disease Sequencing Project (ADSP) whole-genome sequencing (WGS) dataset: A-CUHS-CU000208-BL-COL-56227BL1, A-CUHS-CU000406-BL-COL-52870BL1, A-CUHS-CU000779-BL-COL-31428BL1, A-CUHS-CU001010-BL-COL-52679BL1, A-CUHS-CU002031-BL-COL-25771BL1, A-CUHS-CU002707-BL-COL-40848BL1, A-CUHS-CU002997-BL-COL-47280BL1, A-CUHS-CU003023-BL-COL-47464BL1, A-CUHS-CU003090-BL-COL-47998BL1, and A-CUHS-CU003128-BL-COL-49696BL1. For Illumina250 samples, we used 10 male samples from the Thousand Genomes Project: 4 SAS (HG01583, HG03006, HG03742, NA20845), 3 AMR (HG01112, HG01051, HG01565), 2 EUR (HG00096, HG01500), 1 AFR (HG01879). Finally, we selected our long-read (PacBio and ONT) data from the 1000 Genomes Project, HGSVC3 for the following 10 male samples: 4 AFR (HG01890, HG02666, NA19317, NA19347), 3 EAS (HG01596, NA18534, NA18989), 3 EUR (HG00096, HG00268, HG00358). We specifically selected all male samples to ensure that we include the Y chromosome. Additionally, we specifically chose diverse population backgrounds to minimize bias to any specific population. We aligned the short-read data using bwa-mem [60] and the long-read with minimap2 [61, 62]. All samples were originally aligned to GRCh38 and then aligned to the four different references and have at least 30x average coverage (**Supplementary Fig. S1a-d**).

### Reference Genomes

We used the NCBI version of HG19 with no alternate contigs (https://ftp.ncbi.nlm.nih.gov/genomes/archive/old_genbank/Eukaryotes/vertebrates_mammals/Homo_sapiens/ GRCh37.p13/seqs_for_alignment_pipelines/GCA_000001405.14_GRCh37.p13_no_alt_analysis_set.fna.gz) and HG38 without alternate contigs (https://ftp.ncbi.nlm.nih.gov/genomes/all/GCA/000/001/405/GCA_000001405.15_GRCh38/seqs_for_alignment_pipelines.ucsc_ids/GCA_000001405.15_GRCh38_no_alt_analysis_set.fna.gz). We used the 1000 Genomes version from 2015 of HG38 with alternate contigs (https://github.com/igsr/1000Genomes_data_indexes/blob/master/data_collections/1000_genomes_project/RE ADME.1000genomes.GRCh38DH.alignment). Finally, we used version 2.0 of CHM13 (https://s3-us-west-2.amazonaws.com/human-pangenomics/T2T/CHM13/assemblies/analysis_set/chm13v2.0.fa.gz).

### Epigenetic Data

We pulled WGBS data from (https://www.encodeproject.org/experiments/ENCSR042LOG/) Encode for adrenal gland tissue – (ENCFF384LDT, ENCFF288SYU). We then trimmed reads with TrimGalore (`trim_galore --paired $R1 $R2`) [63] and aligned them to CHM13 with Bismark (`bismark $GENOME -1 $R1 -2 $R2 --parallel 2 --ambig_bam --unmapped --ambiguous`) [64]. We used `--ambiguous` because Bismark removes low MAPQ reads by default, but this flag forces it to write out the ambiguous reads. We then merged the ambiguous reads with the confident reads. We aligned ChIP-Seq data for AD and MCI patients (ENCLB308YWZ, ENCLB142GGP H3K4me3 89 AD; ENCLB305RHN, ENCLB555ZFE H3K4me3 90 MCI) from Encode using bowtie2 to HG38 (`bowtie2 -k 10 -q -x $Ref -p 20 -U $Trimmed_fq -S $Sample_sam`) [65] after trimming the reads with TrimGalore (`trim_galore $file`) [63]. Both assays were filtered using `samtools view` for the annotated genes of interest, then we calculated the coverage using `bedtools genomecov` [66]. We then processed the output using our Epigenetics Rmarkdown script (https://github.com/UK-SBCoA-EbbertLab/DarkRegionCamoPaperFigures/blob/main/EpigeneticsFigs/EpigeneticCamoRegions.Rmd).

The HiC data was plotted and assessed using the UCSC Genome Browser (HESC HiC https://genome.ucsc.edu/cgi-bin/hgTables?db=hg38&hgta_group=regulation&hgta_track=hicAndMicroC&hgta_table=h1hescInsitu&hgta_do Schema=describe+table+schema). The Genome Browser only had the data aligned and analyzed in HG38, which is why we used that reference rather than CHM13 for the figure.

All software for this section was run within a singularity container hosted by Sylabs that can be pulled with the following command: singularity pull --arch amd64 library://mewadsworth/dark_region_followup/epigenetics_software_2023_02_22.sif:sh a256.d9ccbfbb1905e73e09a9bc3237fb5cc70cbe1b1be782e4e52573c0446f40c508.

### DarkRegionFinder (DRF)

Our original paper used a series of bash scripts to run the Dark Region Finder. We converted and updated the scripts to a nextflow pipeline [67] that can be found on Github (https://github.com/UK-SBCoA-EbbertLab/Dark_and_Camouflaged_Genes_Pipeline/tree/master). This runs in conjunction with a singularity container hosted by Sylabs that can be pulled with the following command: singularity pull --arch amd64 library://mewadsworth/dark_region_followup/rescue_camo_variants_2024_10_26.sif:sha256.e9d459cc13d8afa 980900a2981ff84c3072939fabeba38207b6cef034dc91a62. We converted our bash scripts to a self-contained automated Nextflow pipeline. We also provide a web application that creates basic plots such as those in Figures 2, 3, and S2. It also allows the user to filter based on the types of genes and percentage camouflaged.

The Dark and Camouflaged Region identifier is made up of six steps as described in our 2019 work [1]. Briefly, we realigned the samples (for both short- and long-read data) to each respective reference. For this work, we included HG19, HG38 with and without alternate contigs, and CHM13. We used bwa-mem (version 0.7.17-r1188) to align short-read data and minimap2 (version 2.26) for the long-read data. In the original pipeline, we used minimap2’s `map-pb` option for both ONT and PacBio data because it performed better [61, 62]. The current pipeline uses an updated version of minimap2 and we used the `map-ont` for ONT data. The second step of the pipline is to run the Dark Region Finder (DRF) (https://github.com/mebbert/DarkRegionFinder) to identify dark-by-depth regions (regions of less than 5x coverage) and dark-by-MapQ regions (90% of reads have a mapping quality of less than 10). Step three combines the output from the individual samples. Step four prepares the gene annotation bed from the gene annotation GTF file. The fifth step creates the final output bed files for the different types of regions. These bed files are publicly available (https://github.com/UK-SBCoA-EbbertLab/DRF_PaperApp_V2/tree/main/data) and leveraged for the aforementioned WebApp. Finally, the sixth step, which creates a masked genome, is used to rescue variants from camouflaged genomic regions but was not used in this study.

### Including supplementary alignments

To analyze the dark regions including supplementary alignments, we removed the `-M` option used in BWA. The `-M` option marks all supplementary reads determined as small by the algorithm as secondary alignments. Since it would skew our results if we didn’t include all supplementary alignments, we removed that option to keep those reads as supplementary. We additionally added an option to exclude only secondary alignments in htslib used in DRF.

Due to massive increases in depth in some areas when including supplementary reads in our dark region analyses, we limited the number of reads loaded at a time to 10,000 reads in HTSJDK (setMaxReadsToAccumulatePerLocus(10000)). This allowed us to run the DRF with less than 500 gigabytes of memory. Additionally, we removed the intervals for the run with supplementary reads, because if a supplementary read was in a different interval from its primary alignment, HTSJDK failed. The edited pipeline is located on the following branch: https://github.com/UK-SBCoA-EbbertLab/Dark_and_Camouflaged_Genes_Pipeline/tree/nextflow-supplementary-pipeline.

### Web APP & Rscripts

We developed our web application using plotly’s dash app written in R (https://ebbertlab.com/dark_region_comparison.html). The app is split into three tabs: (a) Primary Alignments Only, (b) Primary + Supplementary Alignments, and (c) Comparison. The Primary Alignments Only tab contains plots and tables calculated from the output from the Dark Region Finder app run excluding secondary and supplementary reads. The Primary + Supplementary Alignments tab contains plots and tables calculated from the output from the Dark Region Finder app run excluding only secondary and including primary and supplementary reads. The Comparison tab contains plots and tables created with data from the previous two tabs to compare the results. Most of the plots in this paper were created in and taken from the app. The packages and versions used to deploy the app are listed at the bottom of the page.

Our web app is hosted on Heroku. It uses the heroku-24 stack, an R buildpack (https://github.com/virtualstaticvoid/heroku-buildpack-r) running version 4.4.2, and a basic dyno. Additional installation information can be found in the init.R file on the app’s github page (https://github.com/UK-SBCoA-EbbertLab/DRF_PaperApp/blob/main/init.R). We use the Github deployment method which allows us to deploy the app directly from github.

### IGV screenshots

The different IGV figures were generated using IGV version 2.11.9 [68]. We chose a representative sample from each platform type: Illumina100 - A-CUHS-CU000208-BL-COL-56227BL1, Illumina250 - HG00096, ONT - HG00096, and PacBio – HG00096. Of note we had three sequencing platforms for the 1KG sample HG00096 (Illumin250, PacBio, and ONT). Therefore, when looking at the IGV screen shots the bottom three alignments are for the same sample. We limit the number of insertions and deletions seen by only allowing IGV to show insertions and deletions that are at least 1000 base pairs.

### *CR1* LCR placement

For figure 5, we used data from Brouwers *et al.* to place the *CR1* LCRs using comparison between the two isoforms with and without LCR1’ [53]. We corroborated the location based on the original papers that identified the repeat in the late 1980’s [54, 69] as well as the structure of the C3B binding domain [55].

### Repeat Analysis

We extracted all the reads that were aligned to the area of the beta-satellite and the HSATII using samtools. We then put all of them on the same strand by extracting all reads aligned to the negative strand (samtools view -f 16) and performed a reverse complement using python and then appended them to the forward strand reads (samtools view -F 20). We used RepeatMasker (version 4.1.5) [70] to analyze the repeat structure of the highly variable region on Chromosome 10. First, we analyzed the reads that were completely contained (i.e., did not exceed the boundaries) within the repetitive region (chr10:42,530,255-42,588,899) using the “-s” flag in RepeatMasker [70] to perform a more sensitive search within the curated Dfam library (version 3.7). These results were manually analyzed to identify the repeats in each read within the region. This process was then repeated with all the reads that overlapped with this region to determine if these reads also supported the repeat structure that we identified. RepeatMasker [70] identified the HSATII repeat as alternating “HSATII” and “(CATTC)n simple repeats”, which has been noted previously (Altemose, et al. supplementary materials) [9].

### Annotations

We used the CHM13 repetitive element annotation bed file and intersected it with our dark region bed files using bedtools intersect (https://s3-us-west-2.amazonaws.com/human-pangenomics/T2T/CHM13/assemblies/annotation/chm13v2.0_RepeatMasker_4.1.2p1.2022Apr14.bed). Additionally, for our centromeric satellite annotation comparison from CenSat for CHM13 (https://s3-us-west-2.amazonaws.com/human-pangenomics/T2T/CHM13/assemblies/annotation/chm13v2.0_censat_v2.0.bed). To compare our dark regions against regions that are annotated as unique in CHM13 we leveraged the UCSC Genome Browser CHM13 Unique annotation (https://genome.ucsc.edu/cgi-bin/hgTrackUi?hgsid=1725076402_LoDEV1XMCzGoxZTAnVOGyqEAkw4U&db=hub_3671779_hs1&c=chr5&g=hub_3671779_hgUnique). Finally, to compare the differences between HG38 and CHM13 we compared the Cactus alignment from UCSC Genome Browser (https://genome.ucsc.edu/cgi-bin/hgTables?db=hub_3671779_hs1&hgta_group=compGeno&hgta_track=hub_3671779_cactus&hgta_table<u>=</u> <u>hub_3671779_snakeHg38&hgta_doSchema=describe+table+schema</u>).

### Gene Annotations Used

We used the following gene annotations for each reference.

- HG19 - https://ftp.ensembl.org/pub/grch37/release-107/gff3/homo_sapiens/Homo_sapiens.GRCh37.87.chr.gff3.gz
- HG38 with or without alternates - https://ftp.ensembl.org/pub/release-107/gff3/homo_sapiens/Homo_sapiens.GRCh38.107.chr.gff3.gz
- CHM13 - https://s3-us-west-2.amazonaws.com/human-pangenomics/T2T/CHM13/assemblies/annotation/chm13.draft_v2.0.gene_annotation.gff3

## Contributions

MW and ME developed and designed the study. MW, ME, and MP wrote the manuscript. MW designed and developed the website. MW and MP developed the nextflow pipeline with assistance from BAH. MW, MP, and CS performed all analyses. CS analyzed the chr10 satellite region and advised on the writing of that section. MW developed the DashBio app, and MP embedded the DashBio app into ebbertlab.com. JM provided important intellectual contributions.

## Competing interests

The authors report no competing interests.

## Data availability

Results from our analyses can be found at the following link: https://github.com/UK-SBCoA-EbbertLab/DRF_PaperApp_V2/tree/main. Main results data are available in the data directory in the Github repository. The subfolders labelled “Updated_output_01_17_2025” contain the primary + supplementary alignments. The others contain the primary only alignments.

## Funding

This work was supported by the National Institutes of Health [R35GM138636, R01AG068331 to M.E.], the BrightFocus Foundation [A2020161S to M.E.], Alzheimer’s Association [2019-AARG-644082 to M.E.], PhRMA Foundation [RSGTMT17 to M.E. and predoctoral fellowship to B.A.H].

## Acknowledgments

We appreciate the contributions of the Sanders-Brown Center on Aging at the University of Kentucky. We would like to thank the University of Kentucky Center for Computational Sciences and Information Technology Services Research Computing for their support and use of the Morgan Compute Cluster and associated research computing resources.

We further appreciate and acknowledge data used by both the Alzheimer’s Disease Sequencing Project (ADSP) and the 1000 Genomes Project, and especially the participants of these studies who made this research possible.

The Alzheimer’s Disease Sequencing Project (ADSP) is comprised of two Alzheimer’s Disease (AD) genetics consortia and three National Human Genome Research Institute (NHGRI) funded Large Scale Sequencing and Analysis Centers (LSAC). The two AD genetics consortia are the Alzheimer’s Disease Genetics Consortium (ADGC) funded by NIA (U01 AG032984), and the Cohorts for Heart and Aging Research in Genomic Epidemiology (CHARGE) funded by NIA (R01 AG033193), the National Heart, Lung, and Blood Institute (NHLBI), other National Institute of Health (NIH) institutes and other foreign governmental and non-governmental organizations. The Discovery Phase analysis of sequence data is supported through UF1AG047133 (to Drs. Schellenberg, Farrer, Pericak-Vance, Mayeux, and Haines); U01AG049505 to Dr. Seshadri; U01AG049506 to Dr. Boerwinkle; U01AG049507 to Dr. Wijsman; and U01AG049508 to Dr. Goate and the Discovery Extension Phase analysis is supported through U01AG052411 to Dr. Goate, U01AG052410 to Dr. Pericak-Vance and U01 AG052409 to Drs. Seshadri and Fornage.

Sequencing for the Follow Up Study (FUS) is supported through U01AG057659 (to Drs. PericakVance, Mayeux, and Vardarajan) and U01AG062943 (to Drs. Pericak-Vance and Mayeux). Data generation and harmonization in the Follow-up Phase is supported by U54AG052427 (to Drs. Schellenberg and Wang). The FUS Phase analysis of sequence data is supported through U01AG058589 (to Drs. Destefano, Boerwinkle, De Jager, Fornage, Seshadri, and Wijsman), U01AG058654 (to Drs. Haines, Bush, Farrer, Martin, and Pericak-Vance), U01AG058635 (to Dr. Goate), RF1AG058066 (to Drs. Haines, Pericak-Vance, and Scott), RF1AG057519 (to Drs. Farrer and Jun), R01AG048927 (to Dr. Farrer), and RF1AG054074 (to Drs. Pericak-Vance and Beecham).

The ADGC cohorts include: Adult Changes in Thought (ACT) (U01 AG006781, U19 AG066567), the Alzheimer’s Disease Research Centers (ADRC) (P30 AG062429, P30 AG066468, P30 AG062421, P30 AG066509, P30 AG066514, P30 AG066530, P30 AG066507, P30 AG066444, P30 AG066518, P30 AG066512, P30 AG066462, P30 AG072979, P30 AG072972, P30 AG072976, P30 AG072975, P30 AG072978, P30 AG072977, P30 AG066519, P30 AG062677, P30 AG079280, P30 AG062422, P30 AG066511, P30 AG072946, P30 AG062715, P30 AG072973, P30 AG066506, P30 AG066508, P30 AG066515, P30 AG072947, P30 AG072931, P30 AG066546, P20 AG068024, P20 AG068053, P20 AG068077, P20 AG068082, P30 AG072958, P30 AG072959), the Chicago Health and Aging Project (CHAP) (R01 AG11101, RC4 AG039085, K23 AG030944), Indiana Memory and Aging Study (IMAS) (R01 AG019771), Indianapolis Ibadan (R01 AG009956, P30 AG010133), the Memory and Aging Project (MAP) ( R01 AG17917), Mayo Clinic (MAYO) (R01 AG032990, U01 AG046139, R01 NS080820, RF1 AG051504, P50 AG016574), Mayo Parkinson’s Disease controls (NS039764, NS071674, 5RC2HG005605), University of Miami (R01 AG027944, R01 AG028786, R01 AG019085, IIRG09133827, A2011048), the Multi-Institutional Research in Alzheimer’s Genetic Epidemiology Study (MIRAGE) (R01 AG09029, R01 AG025259), the National Centralized Repository for Alzheimer’s Disease and Related Dementias (NCRAD) (U24 AG021886), the National Institute on Aging Late Onset Alzheimer’s Disease Family Study (NIA-LOAD) (U24 AG056270), the Religious Orders Study (ROS) (P30 AG10161, R01 AG15819), the Texas Alzheimer’s Research and Care Consortium (TARCC) (funded by the Darrell K Royal Texas Alzheimer’s Initiative), Vanderbilt University/Case Western Reserve University (VAN/CWRU) (R01 AG019757, R01 AG021547, R01 AG027944, R01 AG028786, P01 NS026630, and Alzheimer’s Association), the Washington Heights-Inwood Columbia Aging Project (WHICAP) (RF1 AG054023), the University of Washington Families (VA Research Merit Grant, NIA: P50AG005136, R01AG041797, NINDS: R01NS069719), the Columbia University Hispanic Estudio Familiar de Influencia Genetica de Alzheimer (EFIGA) (RF1 AG015473), the University of Toronto (UT) (funded by Wellcome Trust, Medical Research Council, Canadian Institutes of Health Research), and Genetic Differences (GD) (R01 AG007584). The CHARGE cohorts are supported in part by National Heart, Lung, and Blood Institute (NHLBI) infrastructure grant HL105756 (Psaty), RC2HL102419 (Boerwinkle) and the neurology working group is supported by the National Institute on Aging (NIA) R01 grant AG033193.

The CHARGE cohorts participating in the ADSP include the following: Austrian Stroke Prevention Study (ASPS), ASPS-Family study, and the Prospective Dementia Registry-Austria (ASPS/PRODEM-Aus), the Atherosclerosis Risk in Communities (ARIC) Study, the Cardiovascular Health Study (CHS), the Erasmus Rucphen Family Study (ERF), the Framingham Heart Study (FHS), and the Rotterdam Study (RS). ASPS is funded by the Austrian Science Fond (FWF) grant number P20545-P05 and P13180 and the Medical University of Graz. The ASPS-Fam is funded by the Austrian Science Fund (FWF) project I904), the EU Joint Programme – Neurodegenerative Disease Research (JPND) in frame of the BRIDGET project (Austria, Ministry of Science) and the Medical University of Graz and the Steiermärkische Krankenanstalten Gesellschaft. PRODEM-Austria is supported by the Austrian Research Promotion agency (FFG) (Project No. 827462) and by the Austrian National Bank (Anniversary Fund, project 15435. ARIC research is carried out as a collaborative study supported by NHLBI contracts (HHSN268201100005C, HHSN268201100006C, HHSN268201100007C, HHSN268201100008C, HHSN268201100009C, HHSN268201100010C, HHSN268201100011C, and HHSN268201100012C). Neurocognitive data in ARIC is collected by U01 2U01HL096812, 2U01HL096814, 2U01HL096899, 2U01HL096902, 2U01HL096917 from the NIH (NHLBI, NINDS, NIA and NIDCD), and with previous brain MRI examinations funded by R01-HL70825 from the NHLBI. CHS research was supported by contracts HHSN268201200036C, HHSN268200800007C, N01HC55222, N01HC85079, N01HC85080, N01HC85081, N01HC85082, N01HC85083, N01HC85086, and grants U01HL080295 and U01HL130114 from the NHLBI with additional contribution from the National Institute of Neurological Disorders and Stroke (NINDS). Additional support was provided by R01AG023629, R01AG15928, and R01AG20098 from the NIA. FHS research is supported by NHLBI contracts N01-HC-25195 and HHSN268201500001I. This study was also supported by additional grants from the NIA (R01s AG054076, AG049607 and AG033040 and NINDS (R01 NS017950). The ERF study as a part of EUROSPAN (European Special Populations Research Network) was supported by European Commission FP6 STRP grant number 018947 (LSHG-CT-2006-01947) and also received funding from the European Community’s Seventh Framework Programme (FP7/2007-2013)/grant agreement HEALTH-F4-2007-201413 by the European Commission under the programme “Quality of Life and Management of the Living Resources” of 5th Framework Programme (no. QLG2-CT-2002-01254). High-throughput analysis of the ERF data was supported by a joint grant from the Netherlands Organization for Scientific Research and the Russian Foundation for Basic Research (NWO-RFBR 047.017.043). The Rotterdam Study is funded by Erasmus Medical Center and Erasmus University, Rotterdam, the Netherlands Organization for Health Research and Development (ZonMw), the Research Institute for Diseases in the Elderly (RIDE), the Ministry of Education, Culture and Science, the Ministry for Health, Welfare and Sports, the European Commission (DG XII), and the municipality of Rotterdam. Genetic data sets are also supported by the Netherlands Organization of Scientific Research NWO Investments (175.010.2005.011, 911-03-012), the Genetic Laboratory of the Department of Internal Medicine, Erasmus MC, the Research Institute for Diseases in the Elderly (014-93-015; RIDE2), and the Netherlands Genomics Initiative (NGI)/Netherlands Organization for Scientific Research (NWO) Netherlands Consortium for Healthy Aging (NCHA), project 050-060-810. All studies are grateful to their participants, faculty and staff. The content of these manuscripts is solely the responsibility of the authors and does not necessarily represent the official views of the National Institutes of Health or the U.S. Department of Health and Human Services.

The FUS cohorts include: the Alzheimer’s Disease Research Centers (ADRC) (P30 AG062429, P30 AG066468, P30 AG062421, P30 AG066509, P30 AG066514, P30 AG066530, P30 AG066507, P30 AG066444, P30 AG066518, P30 AG066512, P30 AG066462, P30 AG072979, P30 AG072972, P30 AG072976, P30 AG072975, P30 AG072978, P30 AG072977, P30 AG066519, P30 AG062677, P30 AG079280, P30 AG062422, P30 AG066511, P30 AG072946, P30 AG062715, P30 AG072973, P30 AG066506, P30 AG066508, P30 AG066515, P30 AG072947, P30 AG072931, P30 AG066546, P20 AG068024, P20 AG068053, P20 AG068077, P20 AG068082, P30 AG072958, P30 AG072959), Alzheimer’s Disease Neuroimaging Initiative (ADNI) (U19AG024904), Amish Protective Variant Study (RF1AG058066), Cache County Study (R01AG11380, R01AG031272, R01AG21136, RF1AG054052), Case Western Reserve University Brain Bank (CWRUBB) (P50AG008012), Case Western Reserve University Rapid Decline (CWRURD) (RF1AG058267, NU38CK000480), CubanAmerican Alzheimer’s Disease Initiative (CuAADI) (3U01AG052410), Estudio Familiar de Influencia Genetica en Alzheimer (EFIGA) (5R37AG015473, RF1AG015473, R56AG051876), Genetic and Environmental Risk Factors for Alzheimer Disease Among African Americans Study (GenerAAtions) (2R01AG09029, R01AG025259, 2R01AG048927), Gwangju Alzheimer and Related Dementias Study (GARD) (U01AG062602), Hillblom Aging Network (2014-A-004-NET, R01AG032289, R01AG048234), Hussman Institute for Human Genomics Brain Bank (HIHGBB) (R01AG027944, Alzheimer’s Association “Identification of Rare Variants in Alzheimer Disease”), Ibadan Study of Aging (IBADAN) (5R01AG009956), Longevity Genes Project (LGP) and LonGenity (R01AG042188, R01AG044829, R01AG046949, R01AG057909, R01AG061155, P30AG038072), Mexican Health and Aging Study (MHAS) (R01AG018016), Multi-Institutional Research in Alzheimer’s Genetic Epidemiology (MIRAGE) (2R01AG09029, R01AG025259, 2R01AG048927), Northern Manhattan Study (NOMAS) (R01NS29993), Peru Alzheimer’s Disease Initiative (PeADI) (RF1AG054074), Puerto Rican 1066 (PR1066) (Wellcome Trust (GR066133/GR080002), European Research Council (340755)), Puerto Rican Alzheimer Disease Initiative (PRADI) (RF1AG054074), Reasons for Geographic and Racial Differences in Stroke (REGARDS) (U01NS041588), Research in African American Alzheimer Disease Initiative (REAAADI) (U01AG052410), the Religious Orders Study (ROS) (P30 AG10161, P30 AG72975, R01 AG15819, R01 AG42210), the RUSH Memory and Aging Project (MAP) (R01 AG017917, R01 AG42210Stanford Extreme Phenotypes in AD (R01AG060747), University of Miami Brain Endowment Bank (MBB), University of Miami/Case Western/North Carolina A&T African American (UM/CASE/NCAT) (U01AG052410, R01AG028786), and Wisconsin Registry for Alzheimer’s Prevention (WRAP) (R01AG027161 and R01AG054047).

The four LSACs are: the Human Genome Sequencing Center at the Baylor College of Medicine (U54 HG003273), the Broad Institute Genome Center (U54HG003067), The American Genome Center at the Uniformed Services University of the Health Sciences (U01AG057659), and the Washington University Genome Institute (U54HG003079). Genotyping and sequencing for the ADSP FUS is also conducted at John P. Hussman Institute for Human Genomics (HIHG) Center for Genome Technology (CGT).

Biological samples and associated phenotypic data used in primary data analyses were stored at Study Investigators institutions, and at the National Centralized Repository for Alzheimer’s Disease and Related Dementias (NCRAD, U24AG021886) at Indiana University funded by NIA. Associated Phenotypic Data used in primary and secondary data analyses were provided by Study Investigators, the NIA funded Alzheimer’s Disease Centers (ADCs), and the National Alzheimer’s Coordinating Center (NACC, U24AG072122) and the National Institute on Aging Genetics of Alzheimer’s Disease Data Storage Site (NIAGADS, U24AG041689) at the University of Pennsylvania, funded by NIA. Harmonized phenotypes were provided by the ADSP Phenotype Harmonization Consortium (ADSP-PHC), funded by NIA (U24 AG074855, U01 AG068057 and R01 AG059716) and Ultrascale Machine Learning to Empower Discovery in Alzheimer’s Disease Biobanks (AI4AD, U01 AG068057). This research was supported in part by the Intramural Research Program of the National Institutes of health, National Library of Medicine. Contributors to the Genetic Analysis Data included Study Investigators on projects that were individually funded by NIA, and other NIH institutes, and by private U.S. organizations, or foreign governmental or nongovernmental organizations.

## Notes

https://ebbertlab.com/dark_region_comparison.html

### Competing Interest Statement

The authors have declared no competing interest.

