## Supplementary Figures S1-4 for "Sequencing the gaps: dark genomic regions persist in CHM13 despite long-read advances"

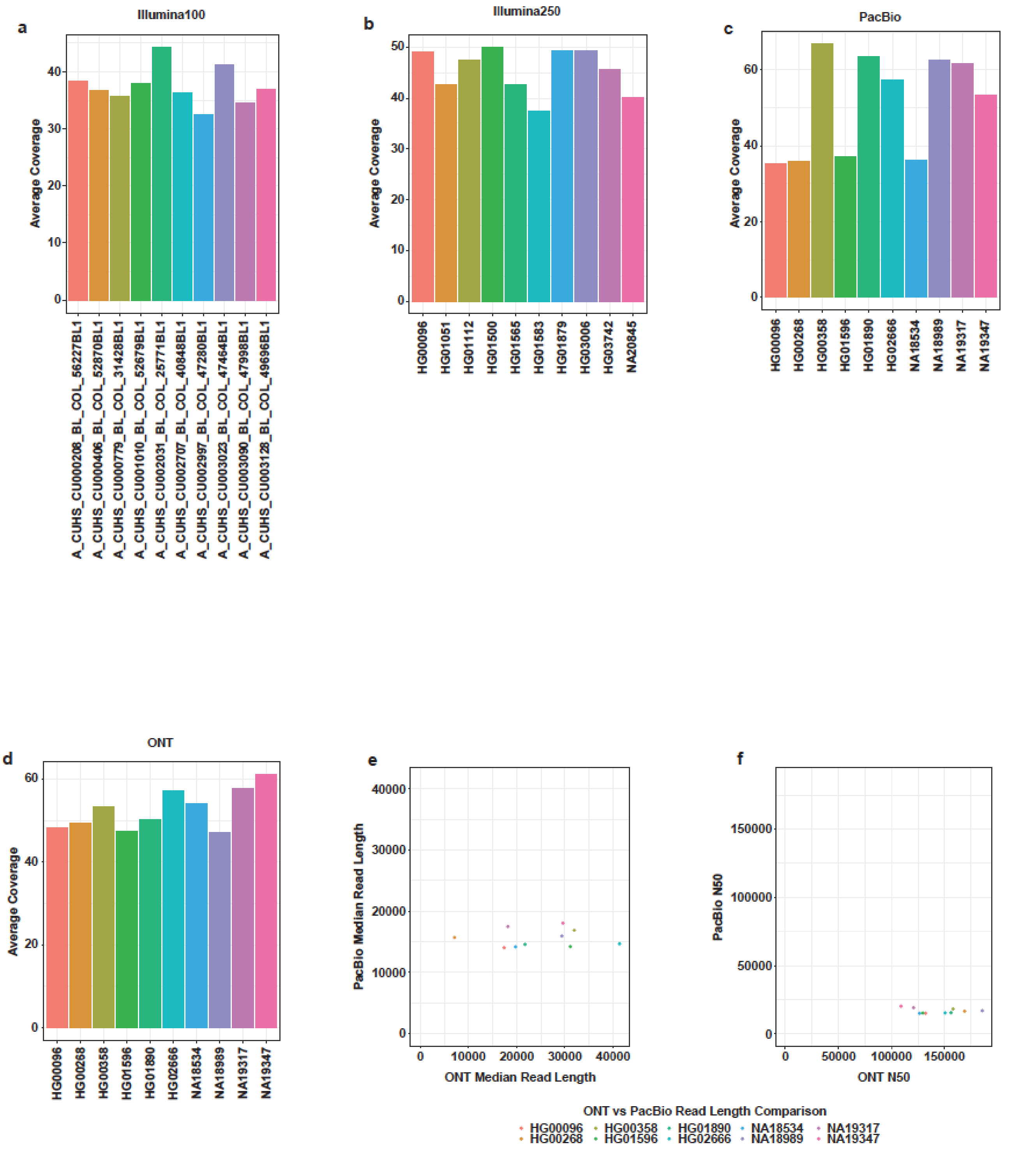


**Figure S1: Sample coverage by platform shows all samples have greater than 30x coverage. (a)** 100 bp Illumina samples all had more than 30x average coverage. **(b)** 250 bp Illumina samples had more than 35x average coverage. **(c)** PacBio HIFI samples also had more than 35x average coverage. **(d)** ONT samples had more than 45x average coverage. **(e)** The median read length of ONT data was mostly much higher than PacBio data. **(f)** The N50 for ONT samples was much higher than PacBio.

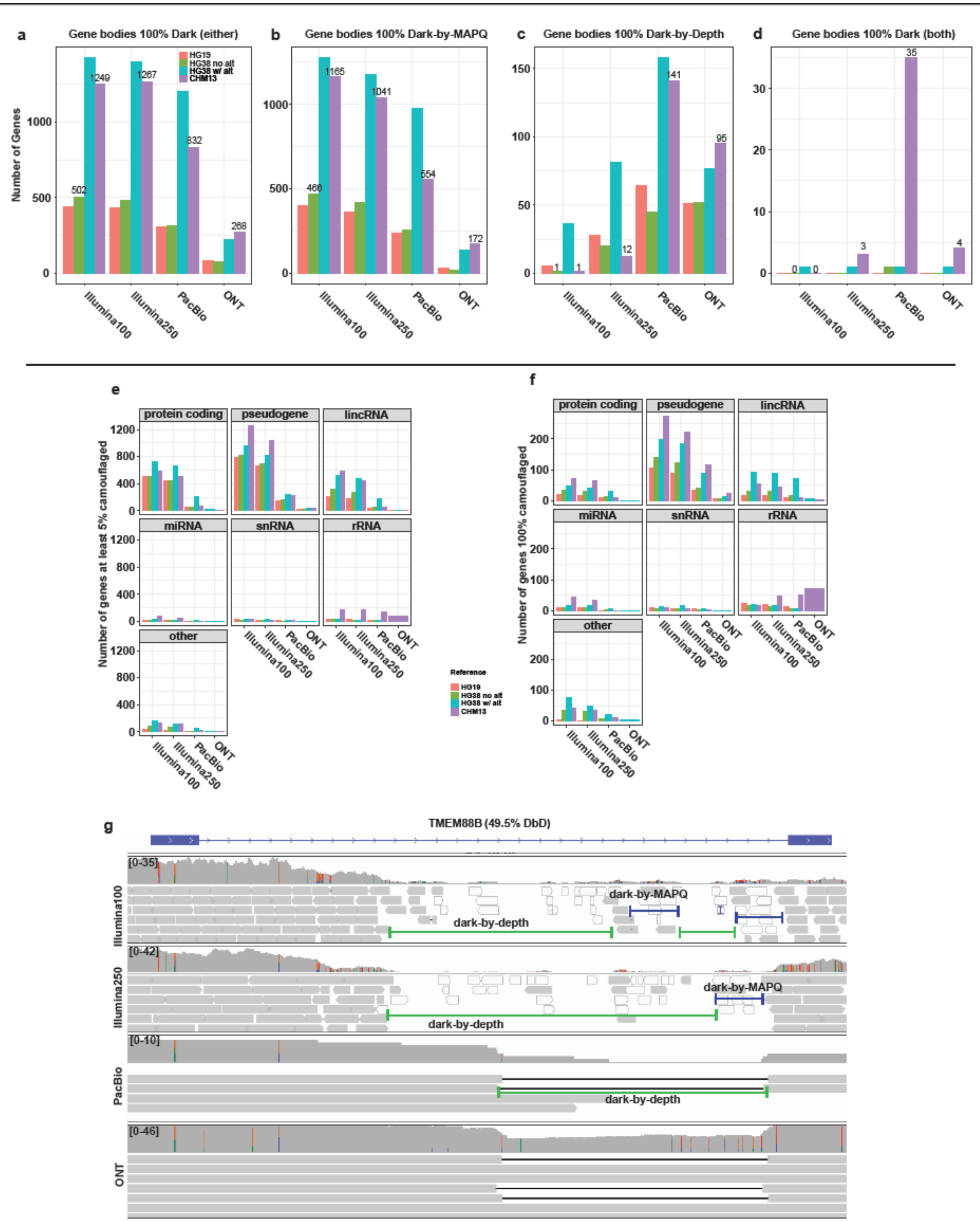


**Figure S2: (a)** Illumina100 and Illumina250 data have the highest number of 100% dark genes followed by PacBio, and then ONT as the lowest. This trend is perpetuated when looking just at genes that are 100% dark-by-depth **(b)** and 100% dark-by-MAPQ **(c)**. **(d)** Surprisingly, we identified a variety of genes that were 100% dark-by-depth and dark-by-MAPQ at the same time. The reasoning here is that, in CHM13, there happen to be multiple annotations for each of the genes in question. In one annotated region it may be dark-by-MAPQ and in another dark-by-depth and therefore come out as 100% of both types of dark region. **(e)** Almost all of the genes that are at least 5% camouflaged are pseudogenes, protein coding, and lincRNA. **(f)** Most of the genes that are 100% camouflaged are pseudogenes. **(g)** TMEM88B which is 49.5% dark-by-depth in Illumina100 data, exhibits both dark-by-depth and dark-by-MAPQ. Long-read data shows that this region is dark because of a heterozygous deletion in the only intron of the gene showing the benefits of using long-read sequencing platforms.

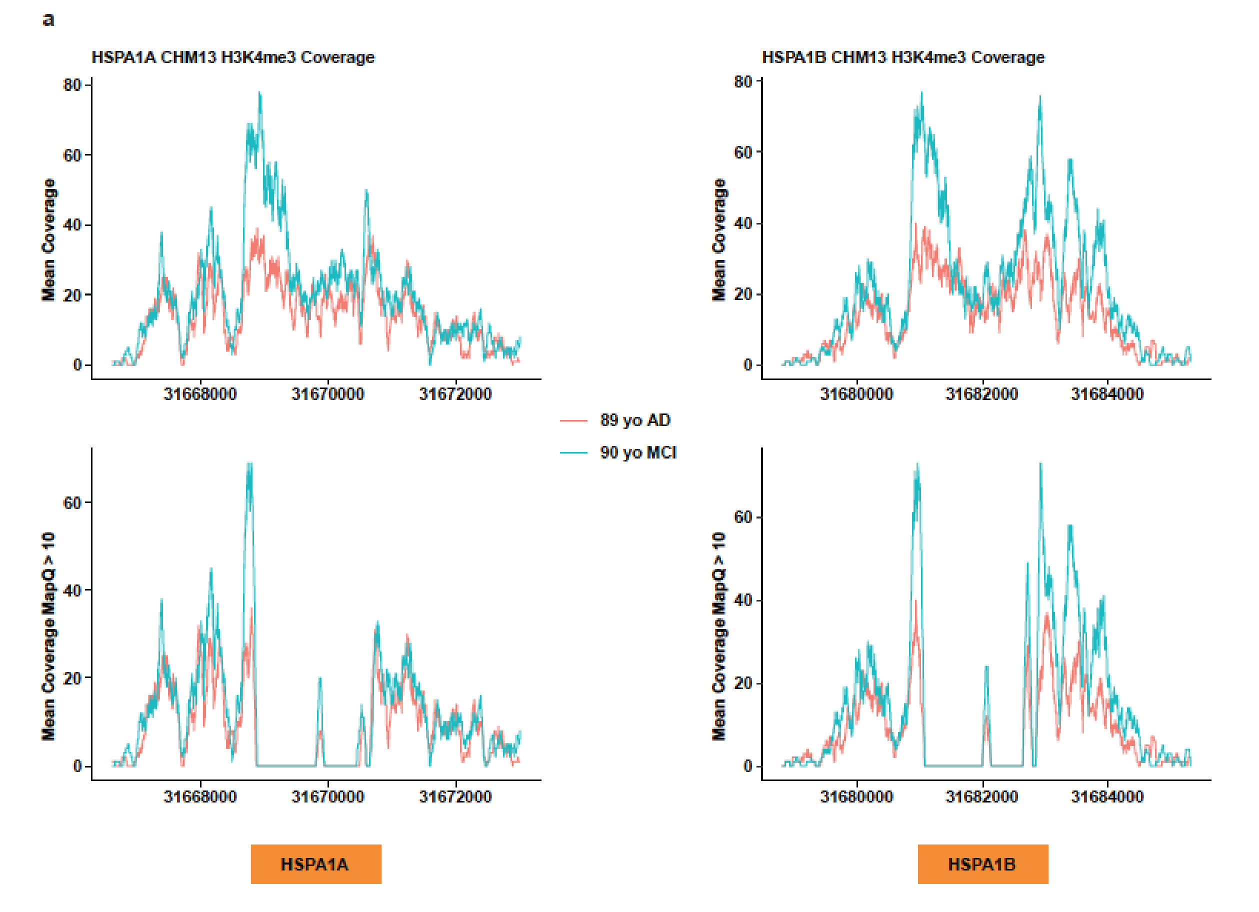


**Figure S3: HSPA1A/B promoter histone H3K4me3 peaks are truncated in ChIP-Seq data. (a)** HSPA1A and HSPA1B are camouflaged to each other and as a result H3K4me3 peaks become truncated and would therefore have either decreased magnitude or be completely ignored during a ChIP-Seq analysis.

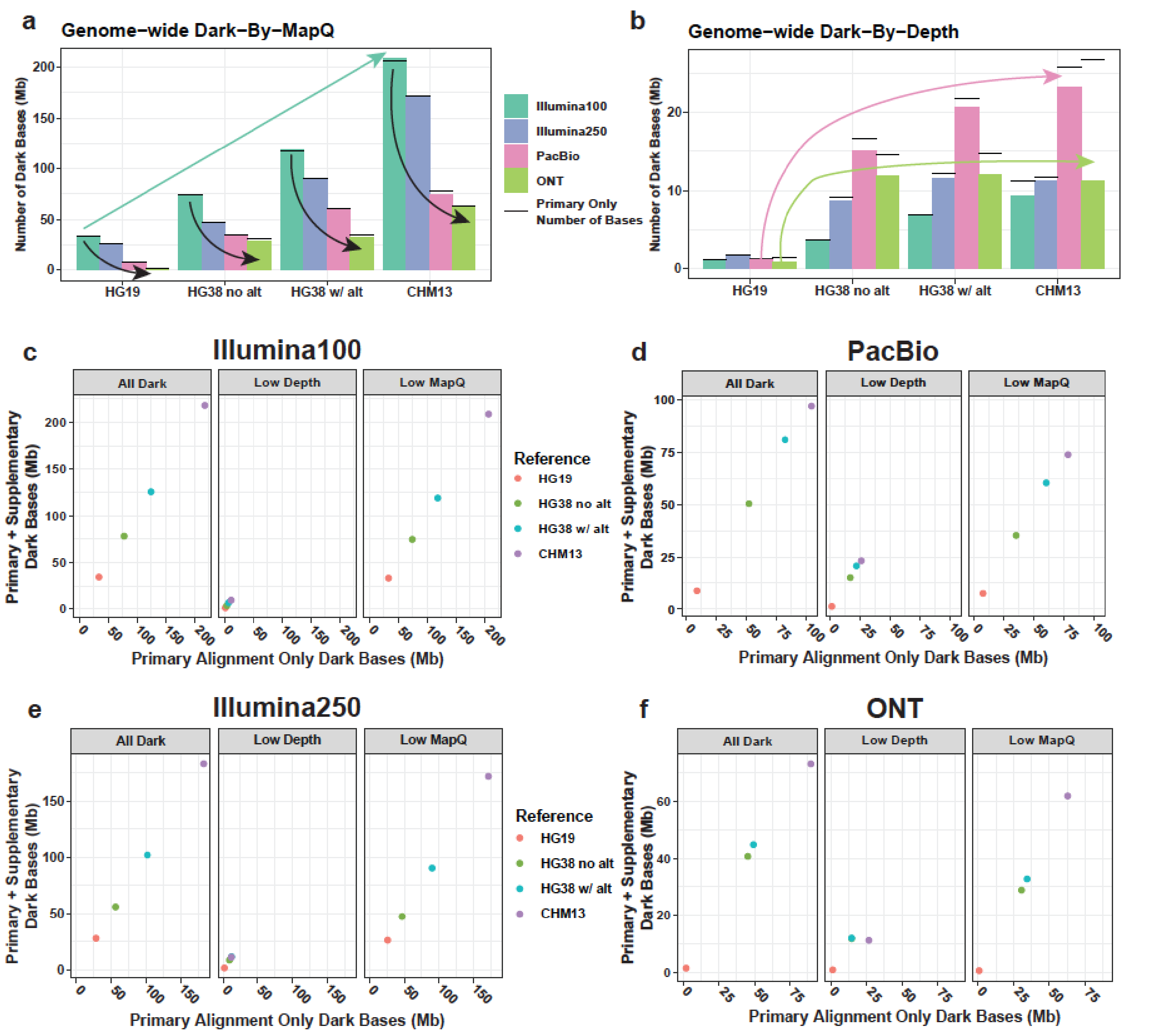


**Figure S4: Including supplementary reads resolves many dark regions. (a)** Genome-wide dark-by-MAPQ bases increase commensurate to the completeness of the genome. Within each reference the longer the reads the less the number of dark bases. The black lines are the levels for primary only alignments and show only marginal changes. **(b)** Genome-wide dark-by-depth bases are the lowest in HG19 and are of similar levels in HG38 with and without alternates and CHM13. Supplementary reads decrease dark-by-depth bases and ONT in CHM13 has the highest number of dark bases resolved. **(c)** Illumina100 dark bases are very similar regardless of the inclusion of supplementary reads. **(d)** PacBio dark-by-depth bases are very similar and are the highest of any sequencing platform regardless of supplementary read inclusion. **(e)** Illumina250 dark bases are very similar regardless of the inclusion of supplementary reads. **(f)** ONT dark-by-depth bases in CHM13 are lower than those in HG38 with supplementary reads are included and does not increase with the completeness of the genome, hinting at structural variation differences in the CHM13 that are resolved by supplementary reads.
